# Single-cell transcriptional landscapes of *Aedes aegypti* midgut and fat body after a bloodmeal

**DOI:** 10.1101/2024.11.08.622039

**Authors:** Thomas Vial, Hélène Lopez-Maestre, Elodie Couderc, Silvain Pinaud, Virginia Howick, Jewelna Akorli, Mara Lawniczak, Guillaume Marti, Sarah Hélène Merkling

**Affiliations:** Institut Pasteur, Université Paris Cité, CNRS UMR2000, Insect Infection & Immunity group, Insect-Virus Interactions Unit, Paris, France; Institut Pasteur, Université Paris Cité, Bioinformatics and Biostatistics Hub, Paris, France; Sorbonne Université, Collège Doctoral, 75005 Paris, France; MIVEGEC, Université de Montpellier, IRD, CNRS, Montpellier, France; School of Biodiversity, One Health and Veterinary Medicine, Wellcome Centre for Integrative Parasitology, University of Glasgow, UK; Department of Parasitology, Noguchi Memorial Institute for Medical Research, University of Ghana, P.O. Box LG 581, Legon, Accra, Ghana; Wellcome Sanger Institute, Hinxton, CB10 1SA, UK; Metatoul-AgromiX Platform, LRSV, Université de Toulouse, CNRS, UT3, INP, Toulouse, France; MetaboHUB-MetaToul, National Infrastructure of Metabolomics and Fluxomics, Toulouse, France

## Abstract

*Aedes aegypti* mosquitoes are vectors for numerous arboviruses that have an increasingly substantial global health burden. Following a bloodmeal, mosquitoes experience significant physiological changes, primarily orchestrated by the midgut and fat body tissues. These changes begin with digestion and culminate in egg production. However, our understanding of those key processes at the cellular and molecular level remains limited. We have created a comprehensive cell atlas of the mosquito midgut and fat body by employing single-cell RNA sequencing and metabolomics techniques. This atlas unveils the dynamic cellular composition and metabolic adaptations that occur following a bloodmeal. Our analyses reveal highly diverse cell populations, specialized in digestion, metabolism, immunity, and reproduction. While the midgut primarily comprises enterocytes, enteroendocrine and intestinal stem cells, the fat body consists not only of trophocytes and oenocytes, but also harbors a substantial hemocyte population and a newly found fat body-yolk cell population. The fat body exhibits a complex cellular and metabolomic profile and exerts a central role in coordinating immune and metabolic processes. Additionally, an insect-specific virus, PCLV (Phasi Charoen-Like Virus) was detected in single cells, mainly in the midgut a week after the bloodmeal. These findings highlight the complexity of the mosquito’s abdominal tissues, and pave the way towards the development of exquisitely refined vector control strategies consisting of genetically targeting specific cell populations and metabolic pathways necessary for egg development after a bloodmeal.

## INTRODUCTION

Blood-feeding female *Aedes aegypti* mosquitoes transmit numerous arboviruses (arthropod-borne viruses) including dengue, Zika, chikungunya and yellow fever viruses. These viruses pose a substantial global health risk that is set to be exacerbated by the changing climate and increased urbanization^1,2^. Developing effective control strategies against insect vectors entails a deep understanding of their physiology at the cellular and molecular level. Recent advances in insect genetic manipulation now enable more precise studies and targeting of these vectors, elevating our capacity to devise effective control strategies^3,4^.

When a mosquito bites a vertebrate host to take a bloodmeal, physiological and metabolic changes unfold within the mosquito, orchestrated by two key organs: the midgut and the fat body^5,6^. The midgut digests the bloodmeal, supplying essential nutrients for egg development^5,7^. During this process, the midgut epithelium also undergoes considerable remodeling, including an initial expansion phase followed by a cell-death program to maintain functional digestion capabilities^8^. Upon receiving nutrients from the midgut, the fat body, acting as the metabolic and reproductive hub, engages in vitellogenesis, the production of yolk protein essential for egg development^9^. Operating between 24 and 72 hours after the bloodmeal^10^, this synchronized process is crucial for egg production and species survival. Beyond these primary roles, the midgut and fat body are also pivotal immune organs, serving as the first line of defense against ingested pathogens^11,12^. The midgut epithelium locally produces antimicrobial peptides (AMPs) and reactive oxygen species (ROS)^13^ while the fat body orchestrates systemic humoral and cellular immune responses, involving secreted immune factors (AMPs, cytokines) and activation of hemocytes, the circulating immune cells^14^.

While the mosquito midgut has been well studied on its own^15–17^, limited data considering its spatial proximity with the fat body and potential inter-organ complementarity is available. In addition, the fat body organ has never been studied at single-cell resolution and its two main cell types have been identified using morphological criteria^12,18^. In this study, we exploit the power of single-cell RNA sequencing (scRNA-seq) to characterize midgut and fat body cell populations. We generated single-cell atlases at two critical stages: two days post bloodmeal, marking the completion of digestion and tapering of vitellogenesis, and seven days post bloodmeal, with the return of tissue homeostasis. Analysis of these datasets reveal the cellular landscape and transcriptional signatures associated with mosquito midgut and fat body organs in a comprehensive and integrated manner, demonstrating their complementarity and unveiling specialized roles for very diverse, and sometimes newly described cell populations. We validate key findings by metabolomic profiling, confirming distinct physiological and metabolic states and roles of each organ. Additionally, we detected the presence of the insect-specific virus PCLV (Phasi Charoen-like Virus) in midgut and abdominal fat body tissues at the single-cell level, demonstrating the power of single-cell transcriptomics for characterizing mosquito viromes at the tissular and cellular levels. Finally, we release our transcriptomes via the interactive *Aedes cell Atlas* (https://aedes-cell-atlas.pasteur.cloud), providing a community platform for gene exploration and analysis in *Ae. aegypti*.

## RESULTS

### Mapping the cellular landscape of mosquito midgut and abdominal fat body following a bloodmeal

To examine how a bloodmeal shapes cell populations and gene expression in the mosquito midgut and abdominal fat body, we performed single-cell RNA sequencing (scRNAseq) on female *Ae. aegypti*. We collected midgut and fat body tissues from a field-derived colony originating from Kumasi (Ghana) at 2- and 7-days post bloodmeal (dpbm)^19,20^. For each timepoint, we collected 7 midguts and 7 fat bodies, with each pair originating from one individual. Specifically, after the midgut was collected, the abdomen of each mosquito was dissected to isolate what we refer to as the ‘fat body’, which includes the abdominal fat body and associated tissues (e.g. muscles, epidermis), while excluding all other visible organs (cf. Methods section; **Fig. 1A**). The tissues were then dissociated into single cells using an enzymatic digestion at 4°C^21,22^. Single-cell transcriptomes were generated using the 10X Genomics^©^ technology (cf. Methods section; **Fig. 1A**). After quality control (cf. Methods and **Fig. S1A, C**), we retained 7,512 and 6,935 cells from the midgut at 2 and 7 dpbm, respectively, and 13,400 and 14,326 cells from the fat body at 2 and 7 dpbm, respectively (**Fig. S1B, D, Table S1**). To compare expression profiles across time within each organ, we integrated datasets from both timepoints (2 and 7 dpbm) using Seurat^23^. Next, we identified distinct cellular clusters and markers (**Fig. 1B-C, Fig. S1B, D)** and carried out a comprehensive annotation by manual curation of the literature available in *Drosophila*, *Anopheles*, and *Ae. aegypti,* detailed below.

**Figure 1.**
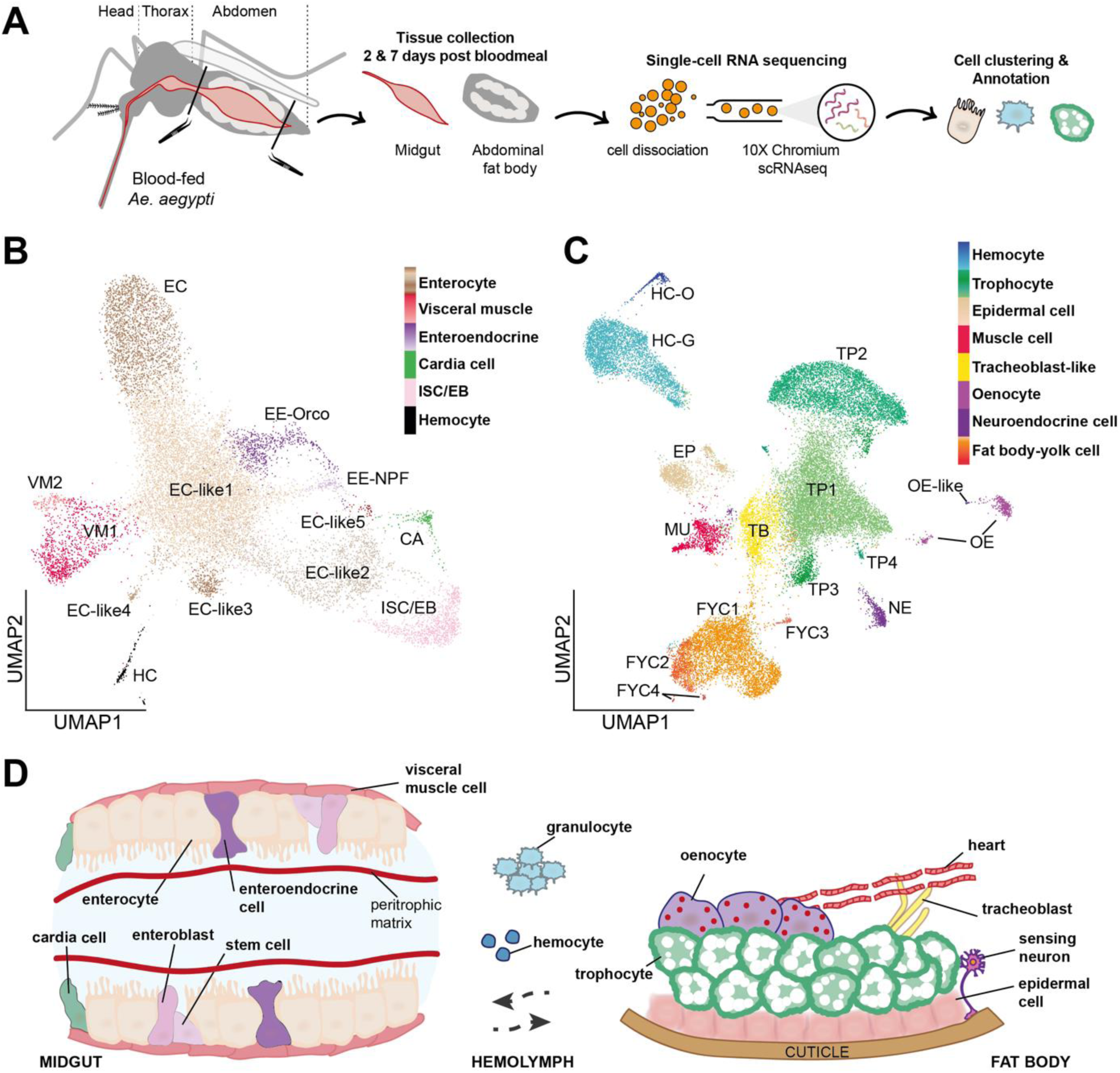
Cellular landscape of *Ae. aegypti* abdominal tissues after a bloodmeal. (**A**) Schematic representation of the experimental workflow. Female *Ae. aegypti* mosquitoes were fed a bloodmeal, and pools of 7 midgut and abdominal fat body tissues were collected at 2 and 7 days post bloodmeal (dpbm), for subsequent cell isolation and single-cell RNA sequencing (scRNAseq). (**B and C**) Integrated UMAP (Uniform Manifold Approximation and Projection) visualization of scRNAseq data from the midgut (**B**) and fat body (**C**) at 2 dpbm and 7 dpbm. Each dot represents a single cell, colored according to its assigned cell cluster. EC: enterocytes; ISC/EB: intestinal stem cells/enteroblasts; EE: enteroendocrine cells; VM: visceral muscle cells; HC: hemocytes; CA: cardia cells; TP: trophocytes; FYC: fat body-yolk cell; HC-G: hemocytes-granulocytes; HC-O: hemocytes-oenocytoids; TB: tracheoblast-like; EP: epidermal cells; MU: muscle cells; NE: neuroendocrine cell; OE: oenocytes. (**D**) Schematic representation of cell types that compose mosquito midgut and fat body associated tissues.

In midguts, we identified 13 cell clusters, encompassing well-known populations such as enterocytes (EC), enteroendocrine cells (EE), intestinal stem cells/enteroblasts (ISC/EB), cardia cells (CA), visceral muscle cells (VM), and hemocytes (HC) (**Fig. 1B**). These main populations were identified by known marker genes, including *nubbin* and serine proteases for enterocytes, *orcokinin* and *prospero* for enteroendocrine cells, *escargot* for intestinal stem cells/enteroblasts, *gambicin* for cardia cells, *titin* for visceral muscle cells, and *NimB2/SPARC* for hemocytes. Our data aligns with established findings from the *Drosophila* midgut single-cell atlas^24^ and previously published single-nuclei and single-cell transcriptomes of *Ae. aegypti* midgut cells^15–17^, validating the robustness of our current scRNAseq approach (**Fig. S2, Table S2**). Additionally, it reveals cellular changes at two critical time points post bloodmeal. The overall cellular composition remained largely stable between 2 and 7 dpbm, with enterocytes representing over 70% of all cells, followed by visceral muscle cells (∼7%) and enteroendocrine cells (∼4-6%) (**Table S2**). We observed a significant increase in intestinal stem cells/enteroblasts at 7 dpbm (3,6% to 11,2%), consistent with a bloodmeal-induced epithelium regeneration^25^.

In fat bodies, we identified 16 cell clusters, revealing a complex cellular ecosystem (**Fig. 1C**). The cellular composition of the fat body was subject to greater temporal changes, demonstrating a high degree of reshuffling after a bloodmeal (**Fig. S1D, Table S3**). Trophocytes (TP), the primary storage sites for blood-derived nutrients, are characterized by the expression of vitellogenic and biosynthetic genes. They constitute the majority of cells at 2 dpbm (59,8% of all cells). However, their proportion significantly declines by 7 dpbm (to 39,3% of all cells), likely reflecting the completion of vitellogenesis and major yolk protein production. This shift in trophocyte abundance over time suggests a return towards a homeostatic state after an intense metabolic activity linked to egg development^26^. Beyond trophocytes, the fat body harbors a diverse range of cell types (**Fig. 1C, Table S3**). First, we detect a population that we named as ‘Fat body-Yolk Cells’ (FYCs; 18% of all cells) expressing a transcriptional profile associated with vitellogenesis and oogenesis, suggesting a link with the physically adjacent ovaries. We also observe a significant increase in *NimB2*/*SPARC*-positive hemocytes at 7 dpbm (18,9% of all cells at day 7, compared to 9% at day 2), likely reflecting heightened immune activity within the fat body post bloodmeal^27^. These are likely sessile hemocytes that associate with tissues in the abdominal cavity^28,29^, and are composed of granulocytes (HC-G) and oenocytoids (HC-O). Tracheoblast-like cells (TB; ∼6%) express genes involved in the regeneration and development of the tracheal system. Muscle cells (MU; ∼3-5%) likely contribute to structural support within the fat body and mosquito heart, while oenocytes (OE;∼1-3%) contribute to lipid synthesis. Epidermal cells (EP; ∼2-7%) and neuroendocrine cells (NE; ∼1-3%) might be involved in barrier function and regulation of various hormonal and physiological processes.

Collectively, we have determined the cellular landscape of midgut and fat body tissues in *Aedes aegypti,* 2 and 7 days after a bloodmeal and propose a sketch of both organs based on our cellular atlases (**Figure 1D).** Our single-cell atlases are accessible and explorable on a dedicated website (https://aedes-cell-atlas.pasteur.cloud).

### Cell population dynamics in the midgut: from bloodmeal digestion to the return to homeostasis

The midgut is key to blood digestion and provides the first line of defense against ingested pathogens^30^. While its cellular composition is increasingly understood, the cell population dynamics following a bloodmeal and over extended periods remain largely unexplored. After identifying the main cell types (**Fig.1**), we analyzed each cell cluster separately and provided a set of highly enriched and sometimes specific new marker genes (**Fig. 2A-B**).

**Figure 2.**
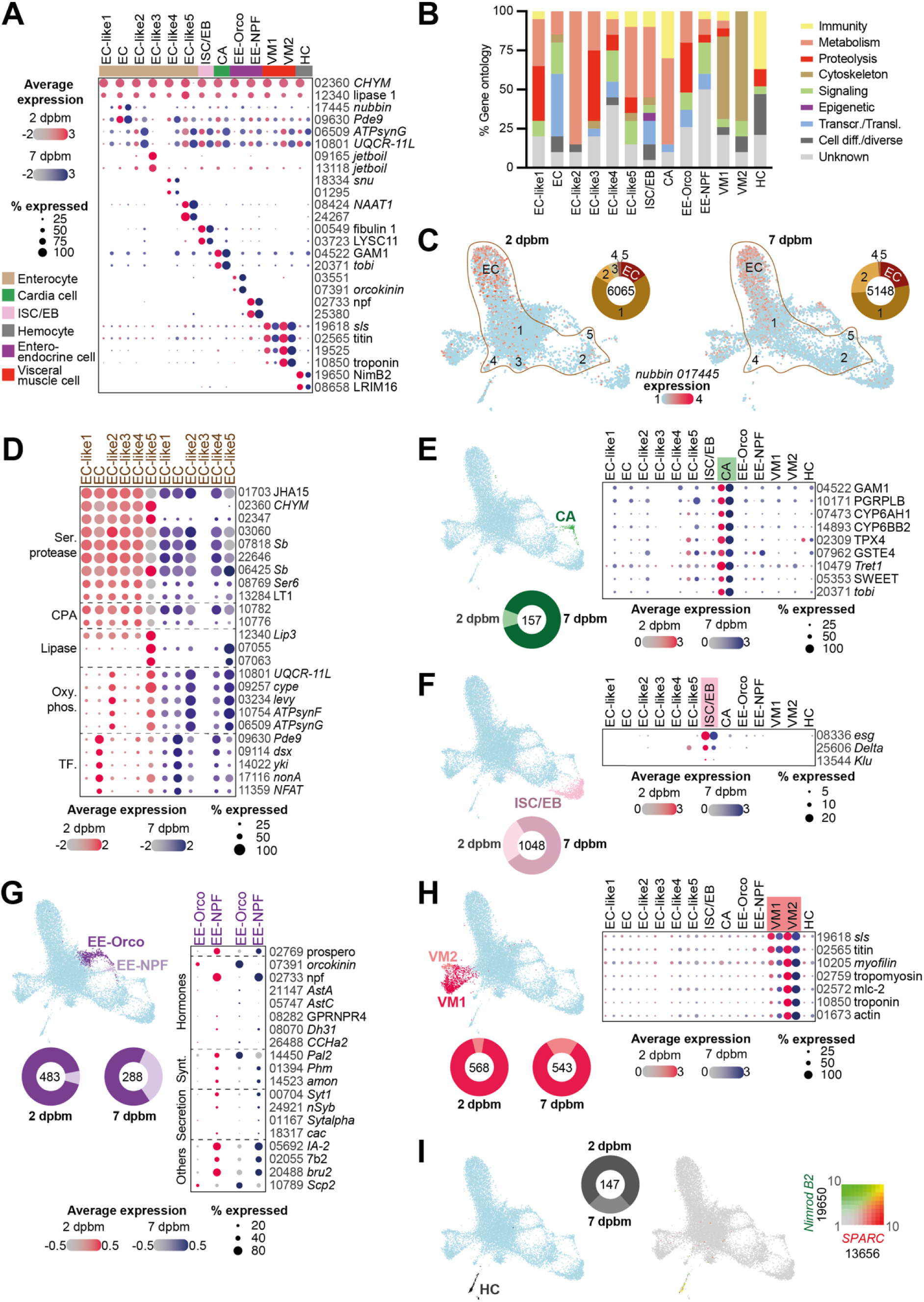
Dynamic midgut cell composition after a bloodmeal. (**A**) Expression of top 2 marker genes for each midgut cell cluster, based on integrated samples collected at 2 and 7 dpbm (Supplementary file 2). (**B**) Gene ontology of top 20 gene markers for each midgut cell cluster. The gene ontology categories were manually curated based on *Aedes aegypti* GO terms, GO terms of *Drosophila* or *Anopheles* orthologs, and annotations of immune-related and digestive-related genes from Hixson *et al*^30^ (Supplementary file 4). Transcr: transcription; Transl: translation; Cell diff: cell differentiation. (**C**) Expression of the *nubbin* (AAEL017445) gene marker in enterocytes (EC) and EC-like clusters at 2 and 7 dpbm, displayed by feature plot. The EC clusters are outlined by a brown line and EC proportion are shown by part of whole plot with total number of cells. (**D**) Dot plot showing the expression patterns of EC and EC-like gene group markers within each cluster at 2 and 7 dpbm. Ser: serine; CPA: carboxypeptidase; Oxy. Phos: oxidative phosphorylation; TF: transcription factor. (**E-H**) Specific cell clusters visualized on UMAP associated with their cell proportions by part of whole plot and gene markers representative of (**E**) cardia cells (CA), (**F**) intestinal stem cells and enteroblasts (ISC/EB), (**G**) enteroendocrine cells (EE) and (**H**) visceral muscle cells (VM). **(I**) Hemocyte cluster (HC) visualized by UMAP and co-expression of *Nimrod B2* (*AAEL019650*) and SPARC (*AAEL013656*) as universal hemocyte gene markers. Dot color shows relative average expression intensity for each gene at 2 dpbm (red) and 7 dpbm (blue). Dot size reflects the percentage of cells expressing corresponding genes in each cell cluster. Each gene is identified by the AAEL0 truncated Vectorbase IDs and their respective abbreviation if available, not italicized. *Drosophila* or *Anopheles* ortholog abbreviations are indicated in italic.

**Enterocytes** (EC). We delineated six distinct enterocyte clusters (EC and EC-like1-5) that are specialized in proteolysis and metabolism (**Fig. 2B**). The cluster we call EC stands out for its enriched expression of *nubbin*, a well-established marker gene for differentiated enterocytes in *Drosophila* and mosquitoes^15,24^ (**Fig. 2A & 2C**). In addition, EC displays enhanced transcriptional and signal transduction activity, with expression of *Pde9*, *doublesex* (*dsx*), and *yorkie* (*yki*), previously linked to *Ae. aegypti* enterocytes^15^ (**Fig. 2D**). Clusters EC-like1-5 likely represent varying functional states of enterocytes and highly express the juvenile hormone-regulated serine protease *JHA15*, previously associated with midgut epithelial cells (**Fig. 2D**) ^24,31,32^. EC-like1 cells express high levels of digestive enzymes (serine proteases, carboxypeptidases, and lipases) at 2 dpbm, indicating robust activity during bloodmeal breakdown^15,24^. EC-like2 and EC-like5 cells express increasing levels of oxidative phosphorylation genes over time, reflecting a transition from digestion at 2 dpbm towards midgut homeostasis by 7 dpbm (**Fig. 2A, D**). Furthermore, the increased abundance of EC- like2 cells at 7 dpbm supports a shift towards midgut stabilization (**Fig. 2D, Table S4**). EC- like3 and EC-like4 cells share digestive markers with EC-like1 at 2 dpbm but exhibit a significant decrease in abundance by 7 dpbm, similar to EC-like1, suggesting these three clusters represent sequential stages or functional states of enterocyte-like cells (**Fig. 2D, Table S4**).

**Cardia cells** (CA). Present mainly at 7 dpbm, cardia cells are characterized by immune and metabolic genes (**Fig. 2A, B, E**). Likely originating from the stomodeal valve near the proventiculus^33^, they express genes encoding antimicrobial peptides such as *gambicin* (*GAM1)*, sugar transporters (*Tret1*, *SWEET*), detoxification and insecticide resistance genes (*TPX4, GSTE4*), and cytochrome P450 genes, potentially contributing to food processing at the juncture with the anterior gut (**Fig. 2E**). Of note, the variable presence of cardia cells may originate from the dissection method, which involves cutting near the proventriculus, potentially impacting the consistent collection of these cells.

**Intestinal stem cells/enteroblasts** (ISC/EB). This cluster is identified using established stem cell markers *fibulin 1* and *C-type lysozyme A* (*LYSC11*)^15^ (**Fig. 2A**). This population also express the *Drosophila* ISC markers *escargot* (*esg*) and *Delta*, along with a lower level of the EB marker *Klumpfuss* (*klu*)^34,35^ (**Fig. 2F**). Subclustering allowed us to fully distinguish stem cells from enteroblasts (**Fig. S3A**). ISC display high *esg* and *Delta* expression, while EB display low *esg* and *Delta* but high *nubbin* expression, indicating a transition stage towards enterocytes prominently at 7 dpbm. Both groups display low *prospero* expression, suggesting limited progression towards enteroendocrine cells.

**Enteroendocrine cells** (EE). Similar to *Drosophila*, EE populations express a set of gut hormones^36^ (**Fig. 2A, G**). The EE-Orco cluster, expressing the *orcokinin* neuropeptide gene linked to egg development, contrasts with the EE-NPF cluster, which expresses *neuropeptide F* (*npf)* and *Islet Antigen-2* (*IA-2*) genes involved in regulating lipid synthesis and insulin metabolism^37–39^. The transcription factor *prospero*, crucial for EE differentiation from ISC, is primarily enriched in the EE-NPF cluster^40^. Furthermore, genes associated with hormone synthesis and secretion are differentially expressed between EE-Orco and EE-NPF, suggesting distinct functional roles (**Fig. 2G**).

**Visceral muscle cells** (VM). Two visceral muscle cell populations (VM-1 and VM-2) are identified based on their expression of structural and regulatory muscle proteins (**Fig. 2A-H**). While both VM express high levels of *sallimus* (*sls*) and *titin* genes, VM-2 displays a broader repertoire of muscle genes (*myofilin, tropomyosin, mlc-2, troponin, actin*), suggesting a more specialized contractile function (**Fig. 2H**).

**Hemocytes** (HC). Lastly, a HC population is primarily identified at 2 dpbm, expressing established hemocyte marker genes such as *Nimrod B2*, *SPARC*, and granulocyte marker *LRIM16*^41^ (**Fig. 2A-I**).

### From metabolism to reproduction and defense: the complex cellular landscape of the fat body

Our study establishes the first comprehensive single-cell atlas of mosquito fat body cells, revealing a remarkably diverse cellular landscape (**Fig. 1C, 3A**). As expected, two major cell types previously described in mosquito fat bodies based on their distinct morphology – trophocytes (TP) and oenocytes (OE) – are readily identified through robust expression of metabolism-related genes^5,18,42^ (**Fig. 3A,B**).

**Figure 3.**
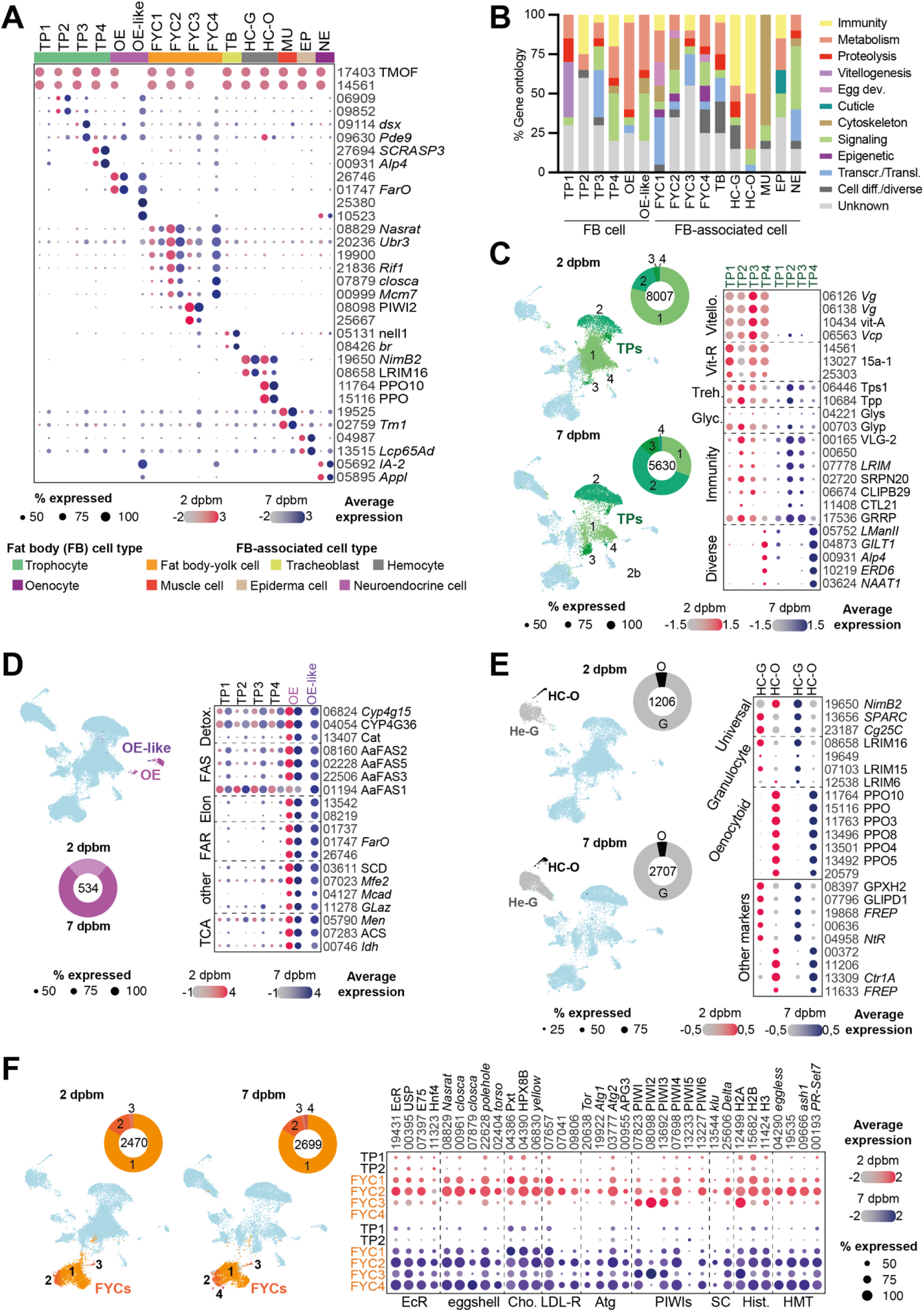
Cellular mosaic of abdominal fat body tissues after a bloodmeal. (**A**) Expression of top 2 marker genes for each fat body cell cluster, based on integrated samples collected at 2 and 7 dpbm (**Supplementary file 3**). (**B**) Gene ontology of top 20 gene markers for each cell cluster. The gene ontology categories were manually curated based on *Aedes aegypti* GO terms, GO terms of *Drosophila* or *Anopheles* orthologs, and annotations **of** immune-related and digestive-related genes from Hixson *et al* (**Supplementary file 4**). Egg dev: egg development; Transcr: transcription; Transl: translation; Cell diff: cell differentiation. (**C-F**) Specific cell clusters visualized on UMAP associated with their cell proportions by part of whole plot and gene markers representative of (**C**) trophocytes, (**D**) oenocytes (**E**) hemocytes and (**F**) fat body-yolk cell. Dot color shows relative average expression intensity for each gene at 2 dpbm (red) and 7 dpbm (blue). Dot size reflects the percentage of cells expressing corresponding genes in each cell cluster. Each gene is identified by the AAEL0 truncated Vectorbase IDs and their respective abbreviation if available, not italicized. *Drosophila* or *Anopheles* ortholog abbreviations are indicated in italic. Vittelo.: vitellogenin genes; Vit-R: vitellogenin membrane receptor; Treh: trehalose metabolic genes; Glyc: glycogen metabolic genes; Detox: detoxification genes; FAS: fatty acid synthase; Elon: elongase; FAR: Fatty acyl-CoA reductase; TCA: tricarboxylic acid cycle genes; EcR: nuclear ecdysone receptor; Cho: chorion; LDL-R: low-density lipoprotein receptor; Atg: autophagy; SC: stem cell gene markers; Histone: histone protein genes; HMT: histone methyltransferase.

**Trophocytes** (TP). Four trophocyte clusters (TP1-TP4) are identified, differentiated by the presence of unique vitellogenic markers and distinct gene expression profiles (**Fig. 3A, C**). TP1 displays a pronounced shift in cellular abundance from 47% at 2 dpbm to 12% at 7 dpbm, suggesting a dynamic remodeling in response to nutrient status changes (**Fig. 3C, Table S5**). TP1 and TP2, representing the majority of trophocytes, show significant expression of vitellogenin genes (*Vg, vit-A, Vcp*) at 2 dpbm, aligning with the peak of proteolytic activity in the midgut associated with yolk protein synthesis^43^ (**Fig. 3B, C**). Beyond vitellogenesis, TP2 cluster, particularly prominent at 7 dpbm, also engages in trehalose biosynthesis, as evidenced by strong expression of *Tps1* and *Tpp* genes. This highlights trophocyte’s crucial role in maintaining energy homeostasis and synthesizing the primary circulatory sugar in insect hemolymph^44,45^. TP2 and TP3 also display a diverse genetic repertoire, encompassing metabolic genes along with a suite of immune modulatory markers (*Vago-like* genes, LRIMs, SERPINs, CLIPs, and lectins). Notably, both clusters expressed the abdomen-specific antimicrobial peptide *holotricin* (*GRRP*)^30^ (**Fig. 3C**). These data highlight the dual role of fat body cells in metabolism and defense, as observed in *Anopheles* mosquitoes^46^. Although TP4 shares the vitellogenin membrane receptor *15a-1* with TP1, it shows a unique signature enriched for lysosomal function (*LManII*), and metabolism/antioxidant defense (*GILT, Alp4*) (**Fig. 3C**). TP3-4 may represent subsets of TP1-2, transitioning towards broader functions beyond yolk production, as indicated by their increasing proportion at 7 dpbm.

**Oenocytes** (OE). Representing only 1-3% of the fat body cell population at both 2 and 7 dpbm (**Fig. 3A, D; Table S5**), oenocytes stand out for their robust metabolic and fatty acid synthesis machinery^47–49^. Specialized in producing cuticular hydrocarbons and pheromones in *Drosophila*^50,51^, they express high levels of genes encoding fatty acid synthases (*AaFAS2, AaFAS3, AaFAS5*), elongase, reductase (*FarO*), desaturase (*SCD*), and P450 enzymes (CYP4G family) (**Fig. 3D**). Notably, oenocytes also exhibit activity in lipid degradation (*Mfe2, Mcad*), transport (*Galz*), and the TCA cycle (*Men, ACS, Idh*), highlighting their multifaceted role in mosquito energy homeostasis. Intriguingly, *AaFAS1* is broadly detected in all fat body cells and overexpressed in the TP2 subpopulation, suggesting distinct functions for AaFAS family members. This is particularly relevant considering *AaFAS1*’s recent identification as a major fatty acid synthase and potential dengue virus proviral factor^52^. Moreover, our findings align with reports of *AaFAS3* and *FarO* as prominent markers of abdominal tissue, significantly contributing to bulk transcriptomic datasets^30^. Additionally, a minor OE-like cluster is detected at 7 dpbm and subclustering revealed a specific gene set associated with this cell population (**Fig. S3B**). Despite their low abundance, oenocytes appear to strongly contribute to the mosquito’s overall metabolism.

**Hemocytes-Granulocytes/Oenocytoids** (HC-G/O). Contrasting with the midgut, the fat body display a diverse landscape of hemocytes, with their abundance doubling at 7 days post-bloodmeal (**Fig. 3E, Table S3**). We identify 2 hemocyte clusters using established universal markers^41^ (*NimB2, SPARC, Cg25C/Col4a1*), likely delineating circulating and sessile populations within the fat body^28,53^ (**Fig 3A, E**). The granulocyte cluster HC-G, express markers of mature granulocytes^41^ (*LRIM16, LRIM15*). In addition, HC-G displays a wide repertoire of stress and immune response genes^30,54^ (*GPXH2, CLIPD1, FREP*), highlighting their potential contribution to local immunity within the fat body. Further subclustering reveals additional heterogeneity within HC-G, ranging from antimicrobial granulocytes, prohemocytes to unidentified granulocytes that differ from circulating hemocytes typically observed in *Ae. aegypti*^46^ (**Fig. S3C**). Particularly, a subcluster highly abundant at 7 dpbm (Gra-Cut), exhibit a distinct profile with lower granulocyte marker expression and a specific increase in cuticle protein (*CPLCA3, CPF3, Ccp84A*) and trypsin (*Sb*) genes (**Fig. S3C**). This suggests a potential specialization of granulocytes towards abdominal/cuticle tissues, particularly at later stages post bloodmeal^28^. Finally, a distinct population (HC-O) express several PPO gene family, a hallmark of oenocytoids with roles in melanization and immune signaling^41,46,55^ (**Fig. 3E, S3C**). These findings suggest a diverse and dynamic hemocyte population associated with or circulating near the fat body after a bloodmeal, likely contributing to local or systemic immune responses induced by the bloodmeal^56^.

**Fat body-yolk Cells** (FYC). We uncovered an undescribed cell population within the fat body of female *Ae. aegypti* following a bloodmeal and named them fat body-yolk cells (FYC) due to their expression of genes essential to fat body and ovary function (**Fig. 3A, B**). FYC uniquely express a suite of genes critical for reproductive processes such as *Nasrat*, crucial for eggshell formation (**Fig. 3A**). Mosquitoes in our experiment received a bloodmeal, without oviposition sites (cf. Methods section). By 7 dpbm, we observed ovaries filled with mature eggs, indicative of an oviposition readiness typically beginning by 3 dpbm^5^. At 2 and 7 dpbm, the FYC population remains stable (**Fig. 3F, Table S3**), suggesting a role in assuring reproductive readiness^5^. Clustering suggests that FYC1, FYC2, and FYC4 likely represent different states of the same cell population, based on shared expression of ecdysone receptor genes (*EcR, USP, E75, Hnf4*) crucial for oogenesis regulation^10,57–59^, and structural proteins (*Nasrat, closca, polehole, torso*) necessary for eggshell formation and embryo protection^60–62^. Additionally, FYC1-2-4 express genes required for eggshell integrity (*Pxt, HPX8B, yellow*)^60^, and genes associated with the LDL receptor family, suggesting a role in ovary lipid transport^63,64^. The enrichment of histone protein genes (*H2A, H2B, H3*) and histone methyltransferase genes (*eggless*, *ash1*, *PR-Set7*) hint at potential epigenetic remodeling activity^65,66^. In addition, FYC highly express autophagy genes (*Tor, Atg1, Atg2, APG3*), suggesting resource reallocation during late vitellogenesis and egg maturation^26,67^. Moreover, FYC2 and FYC4 exhibit higher transcriptional activity and express stem cell markers (*klu, Delta*), suggesting a role in fat body differentiation and regeneration. Interestingly, FYC highly express piRNA pathway (PIWI family) genes that are crucial for transposon silencing and genomic defense in ovarian development ^30,65,68^. FYC3 was particularly enriched in *PIWI2*, hinting to its role in germline protection. Overall, the newly identified FYC population sheds light on *Ae. aegypti* adaptive mechanisms that possibly maintain reproductive readiness and cellular integrity despite environmental restrictions on egg laying^69^. This resilience underscores the sophisticated biological orchestration ensuring the mosquito’s reproductive potential and life cycle continuity.

Four less abundant populations are identified in association with the fat body (**Fig. S4**). **Tracheoblasts** (TB). This cluster displays markers of growth and cell regeneration (*nell1, br, rgn, goe*), as well as signaling and trachea cell marker (*sprouty, Egfr*)^70–72^ (**Fig. 3A, S4A**). In addition, these cells express specific metabolic genes (*Gcat, Tdh*). Their constant presence at 2 and 7 days post bloodmeal suggests a role in tissue maintenance and regeneration. **Muscle cells** (MU). A distinct muscle cell cluster remains stable in proportion between 2 and 7 dpbm, expressing genes characteristic of muscle structure and regulation (**Fig 3A, S4B**). These genes encompassed a variety of muscle fiber types and regulatory pathways shared with cardiac and other muscle cells found in the adult *Ae. aegypti* heart ^73–75^. **Epidermal cells** (EP). A distinct cluster is enriched for epidermal cell markers, with cell abundance increasing at 7 dpbm (**Fig. 3A, S4C**). Epidermal cells express significant levels of juvenile hormone binding protein (*JHbp14*) and genes associated with cuticle formation (*Lcp65Ad, Cpr49Ab, Cpr49Ae*, *dumpy*) crucial for epidermal-cuticle attachment^76^. Cuticular proteins, associated to the most abundant EP subcluster (**Fig. S3D**), are known to be either synthesized by epidermal cells or transported from the hemolymph^77,78^. In addition, epidermal cells display markers for adhesion (*SCRB5, Tsp, CTLGA3*). Interestingly, the tyrosine amino acid pathway appears specifically expressed (*Hpd, hgo, Tat*), which is essential in insect cuticle pigmentation and homeostasis. Lastly, epidermal cells distinctly exhibit expression of acetylcholinesterase (*Ace*) and odorant binding proteins (*OBP11, OBP10*), suggesting their involvement in neuronal signaling and chemosensation. **Neuroendocrine cells** (NE). A NE cell cluster, prominent at 7 dpbm (**Fig 3A, S4D**), is identified based on markers shared with *Drosophila* brain neuropeptides and midgut EE cells (**Fig. 2G**). Neuroendocrine cells notably express amyloid precursor protein (*Appl*) and nervous system antigen (*nrv*), key components in insect neuronal signaling^80,81^. Enriched expression of neuro-specific genes, including *futsch*, a GABA receptor (*Rdl*), and acetylcholinesterase (*Ace*), further emphasizes the broad spectrum of neuronal functions manifested within these cells, underpinning complex neuroendocrine regulation in the fat body. Interestingly, the genes *Ace* and *Rdl*, coding for proteins targeted by insecticides in mosquitoes^82^, highlight the potential role of these cells in environmental adaptation and underscore their significance as precise targets in insecticide-based control strategies.

The abdominal fat body environment of *Ae. aegypti*, traditionally viewed as a metabolic center, is highly complex. Our study unveils distinct trophocyte subpopulations, oenocytes, and a diverse hemocyte population, suggesting multifaceted roles spanning metabolism, immunity and reproduction. Furthermore, the identification of fat body-yolk cells and the presence of epidermal, tracheoblast, muscular, and neuroendocrine cells highlight the fat body as a microenvironment orchestrating diverse physiological processes.

### Specific midgut and fat body cells mutually contribute to immune and metabolic pathways

To examine the respective roles of midgut and abdominal fat body tissues in *Ae. aegypti* following a bloodmeal, we focused on key genes implicated in immune response and metabolism at 2 and 7 dpbm (**Fig. 4**). We computed the proportion of cells expressing specific gene sets per cluster to assess gene expression. The use of control reference genes, including the digestive enzyme *Chymotrypsin-1* (CHYM1) and the vitellogenic protein *Vitellogenin A* (vit-A), confirmed a steady physiological status for each condition. Moreover, the consistent expression of the housekeeping gene 60S ribosomal protein L14 (*RpL14*) across all clusters confirmed overall data quality, with small variations only observed in the midgut at 2dpbm. Both midgut and fat body organs are pivotal in nutrient metabolism and coordination of immune defenses^83–85^. Given these established roles, we decided to analyse expression of canonical immune pathways traditionally associated with host defense, specifically TOLL, IMD, JAK-STAT, antimicrobial peptides and RNA interference.

**Figure 4.**
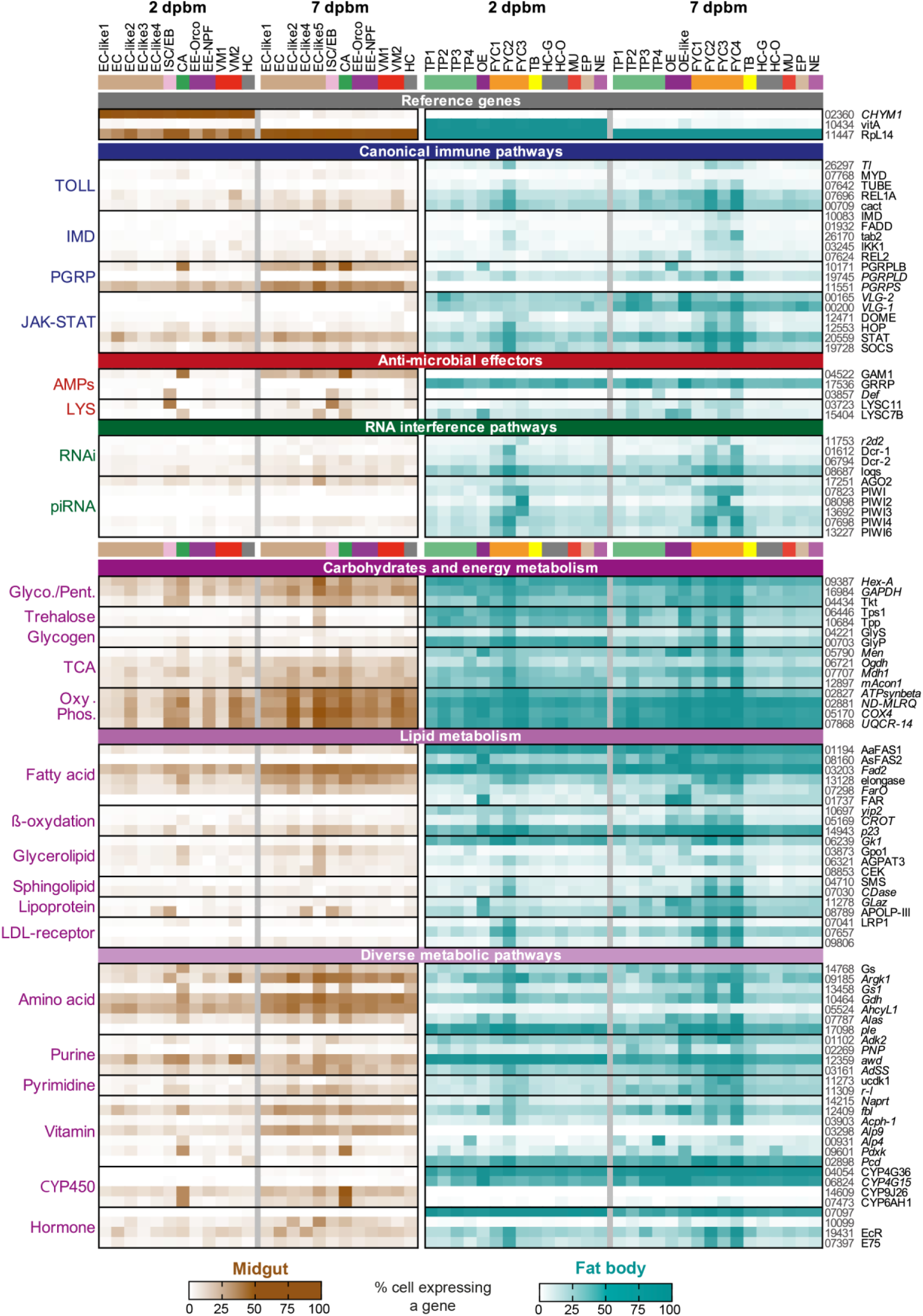
Immune and metabolic pathways are expressed mainly by fat body cells and disparately in midgut cells. The heatmap displays the proportion of cells expressing specific genes within each cluster of midgut and fat body cells. Genes were chosen for their role in key immune and metabolic pathways (left column), with a criterion of at least 15% expression in a minimum of one cluster across the tissues (**Supplementary file 5**). The heatmap features the expression of Chymotrypsin-1 (*CHYM*) and Vitellogenin A (*vitA*) to monitor physiological changes post-bloodmeal, and 60S ribosomal protein L14 (*RpL14*) as a housekeeping gene. Each gene is identified by the AAEL0 truncated Vectorbase IDs and their respective abbreviation if available, not italicized (right column), with *Drosophila* or *Anopheles* ortholog abbreviations in italics. PGRP: PeptidoGlycan Recognition Protein; LYS: lysozyme; Glyco: glycolysis; Pent: pentose phosphate pathway; TCA: tricarboxylic acid cycle; Oxy. Phos: oxidative phosphorylation; CYP450: cytochrome P450.

**TOLL/IMD** pathway components, including *REL1A*, *cactus*, and *REL2*, are broadly expressed across cell clusters, and particularly enriched in EC, CA, VM, and FYC clusters. IMD signaling is specifically upregulated in FYC at 7 dpbm. Peptidoglycan recognition proteins (PGRP), key activators of TOLL/IMD signaling, exhibit tissue-specific expression patterns^86^. *PGRP-LB* is predominant in midgut cells (EC, CA) and OE, while *PGRP-LD* is enriched in the fat body (TP, FYC). *PGRPS* is specifically expressed in midgut cells, with increased expression at 7 dpbm. The **JAK-STAT** pathway is predominantly activated in the fat body. Cytokine-like *Vago-like* genes (*VLG-2, VLG-1*) are significantly expressed in TP2-3, with an increase at 7 dpbm, and to a lesser extent in FYC and EP clusters^87^. Core components of this pathway, including *DOME*, *HOP*, and *STAT*, are broadly expressed in FYC. *STAT* is the only member of this pathway highly expressed in midgut cells, particularly by EC. Cell clusters with high *STAT* expression in both tissues also express the *SOCS* regulator at a lower level.

**Antimicrobial peptides (AMPs)** genes are differentially expressed across tissues. *Gambicin* (*GAM1*) is predominantly expressed in midgut CA cluster at 2 dpbm, expanding to the other midgut clusters at 7 dpbm. *Holotricin* (*GRRP*) is extensively expressed across all fat body clusters, particularly in TP2-3. Other antimicrobial peptides, such as *defensin* (*Def*) and lysozymes, display distinct patterns in the midgut (CA and ISC/EB) and the fat body (TP2 and OE). **RNA interference (RNAi)** pathway components are extensively expressed in the fat body, with FYC exhibiting particular activity for key genes such as *Dcr-1, Dcr-2*, *loqs*, and PIWIs. This highlights the crucial role of FYC in preserving the mosquito’s germline integrity.

The midgut and fat body tissues in mosquitoes are highly metabolically active, particularly following a bloodmeal^43,83,88^. This is demonstrated by a high proportion of cells in both tissues expressing a broad array of metabolic genes, exceeding those involved in immune pathways, observed not only at the end of digestion but also at 7 dpbm (**Fig. 4**).

**Carbohydrate metabolism** including glycolysis and the pentose phosphate pathway (*Hex-A, GAPDH, Tkt*), is active in both tissues, with specific cell groups showing elevated activity: EC, ISC/EB, CA in the midgut, TP2-3, OE and FYC in the fat body. Starch metabolism, particularly trehalose synthesis (*Tps1, Tpp*), is predominantly a fat body function, with a dynamic expression pattern. Glycogen metabolism displays distinct profiles, with glycogen catabolism (*GlyP*) highly active in the fat body at 2 dpbm trough TP and FYC clusters and minimal biosynthesis (*GlyS*) in both tissues. **Energy metabolism**, encompassing the TCA cycle and oxidative phosphorylation, is robust in both tissues, with overlapping cell-type expression. Similar to glycolysis, midgut (EC, CA) and fat body clusters (TP, OE, FYC) show distinct expression patterns for specific genes within these pathways. Notably, oxidative phosphorylation became more widespread at 7 dpbm in both tissues.

**Lipid metabolism** exhibits diverse patterns across the midgut and the fat body. Fatty acid biosynthesis, a core lipid metabolic process, shows tissue-specific profiles. While *AaFAS1* is ubiquitously expressed in the fat body and sporadically by few midgut clusters, *AaFAS2* demonstrates a specific expression pattern led by oenocytes. Interestingly, a fatty acid desaturase (*Fad2*), is consistently expressed in both tissues. Genes involved in fatty acid elongation (*elongase*), reduction (*FarO, FAR*), and beta-oxidation, exhibit diverse expression patterns in midgut (EC-like2, CA, ISC/EB, EE-NPF), and fat body clusters (TP2, OE, and FYC). Glycerolipid and sphingolipid metabolism are expressed relatively low in both tissues. However, a glycerol kinase (*Gk1*), involved in triglycerides and phospholipids synthesis, is predominantly active in TP and FYC cells. Apolipophorins such as *Glaz* and *apoLp-III*, crucial for lipid transport, display cell-specific expression in both tissues, highlighting vital biological communication. *Glaz*, initially expressed in OE cluster, expands to FYC over time. Meanwhile, *apoLp*-III demonstrates dynamic expression across specific midgut (EC-like, CA, ISC/EB) and fat body clusters (TPs, OE, FYC). Lipoprotein receptors (*LRP1, arrow/yolkless*), mainly expressed in the fat body, likely facilitate lipid uptake, particularly in FYCs.

Beyond lipid metabolism, midgut and fat body cell clusters exhibit diverse metabolic functions. **Amino acid** metabolism is especially prominent in the midgut, with pathways focusing on arginine, glutamine, and glutamate (*Argk1, Gs1, Gs, Gdh*) enriched in EC-like, cardia, and VM2 cells. Similarly, the fat body contributes to amino acid metabolism, primarily in OE and FYCs. Notably, a tyrosine/dopamine synthetic gene (*ple*) is exclusive to the fat body, predominantly in FYC. **Purine and pyrimidine** metabolisms show similar patterns between both tissues. The midgut, especially EC-like2, CA and ISC/EB cells, exhibits high expression of key purine genes (*Adk2, awd, AdSS*), while in the fat body elevated expression is notable in TP2 and FYCs. Notably, intensive expression of nucleoside triphosphate regulatory gene (*awd*) is observed across all fat body clusters. **Vitamin** metabolism exhibits diverse, cell-type specific expression profiles. Nicotinate (*Naprt*), pantothenate (*fbl*), and riboflavin (*Acph-1*) pathways, show varying levels of expression in both tissues, with specific enrichment in certain cell types (EC, EC-like, CA, TP, OE and FYCs). Gene related to thiamine (Alp4, Alp9) are mainly represented in diverse midgut clusters, while vitamin B6 (*Pdxk*) displays high expression levels in cardia cells, TP2 and oenocytoids. Additionally, folate metabolism (Pcd) is actively expressed in the fat body. Cytochrome **P450** metabolism, involved in xenobiotic detoxification and associated with insecticide resistance^89^, showcases distinct tissue-specific and cell-type-specific expression patterns. Notably, two CYP4G family genes are prominently expressed in fat body clusters, primarily OE. Meanwhile, the expression of *CYP9J26* and *CYP6AH1* are enriched in midgut CA cluster, highlighting a clear tissue and cell tropism of P450 enzyme activity. **Insect hormone biosynthesis** genes show high levels of expression with distinct tissue specificity, particularly for two 4-nitrophenyl phosphatase genes, which serve as precursors for juvenile hormone synthesis. Notably, no expression of genes directly associated with ecdysone hormone synthesis are observed. However, ecdysone receptor genes, such as *EcR* and *E75*, are broadly expressed at low levels in midgut EC and EC-like cells, as well as in FYCs of the fat body, as previously illustrated (**Fig. 3F**).

Our findings portray the distinct and key roles played by the midgut and fat body, highlighting their unique contributions to immune and metabolic functions after a bloodmeal. The midgut, with enterocytes, cardia cells, and ISC/EB cells, mounts a focused immune response while actively participating in metabolic processes. Conversely, the fat body and associated cells exhibit a broader role in immunity and metabolism, with trophocytes, oenocytes and fat body-yolk cells emerging as key players. Consistent with recent research linking metabolic processes to immune cell functions, our findings suggest that metabolic activity and immune responses are intertwined after blood ingestion in mosquitoes^90^.

### A metabolomic analysis confirms the transcriptional landscape of midgut and fat body tissues post bloodmeal

To further validate our findings, we carried out an extensive metabolomic analysis to characterize physiological states and find metabolic specializations in the midgut and fat body of blood-fed *Ae. aegypti* mosquitoes. In parallel with the scRNA-seq experiment, abdominal tissues dedicated to metabolite extraction were dissected from mosquitoes belonging to the same cohort at 2 and 7 dpbm. Ultra-high-performance liquid chromatography−high-resolution mass spectrometry (UHPLC−HRMS) analysis of pooled midgut and pooled fat bodies (n=5) generated a comprehensive metabolic profile encompassing 974 metabolites, subsequently analyzed by Principal Component Analysis (PCA), (**Fig. 5A**). PCA validates clear tissue-specific and temporal metabolic distinctions, with samples clustering separately based on tissue type and days post bloodmeal (**Fig. 5B**). Differential metabolite abundance analysis reveals distinct metabolic profiles between midgut and fat body tissues (**Fig. 5C**). At 2- and 7- dpbm, 155 and 149 unique metabolites, respectively, differentiate the two tissues, highlighting significant physiological divergence. Moreover, the midgut exhibits a more pronounced time-dependent metabolic shift, with 222 differentially abundant metabolites between 2 and 7 dpbm, compared to 63 in the fat body.

**Figure 5.**
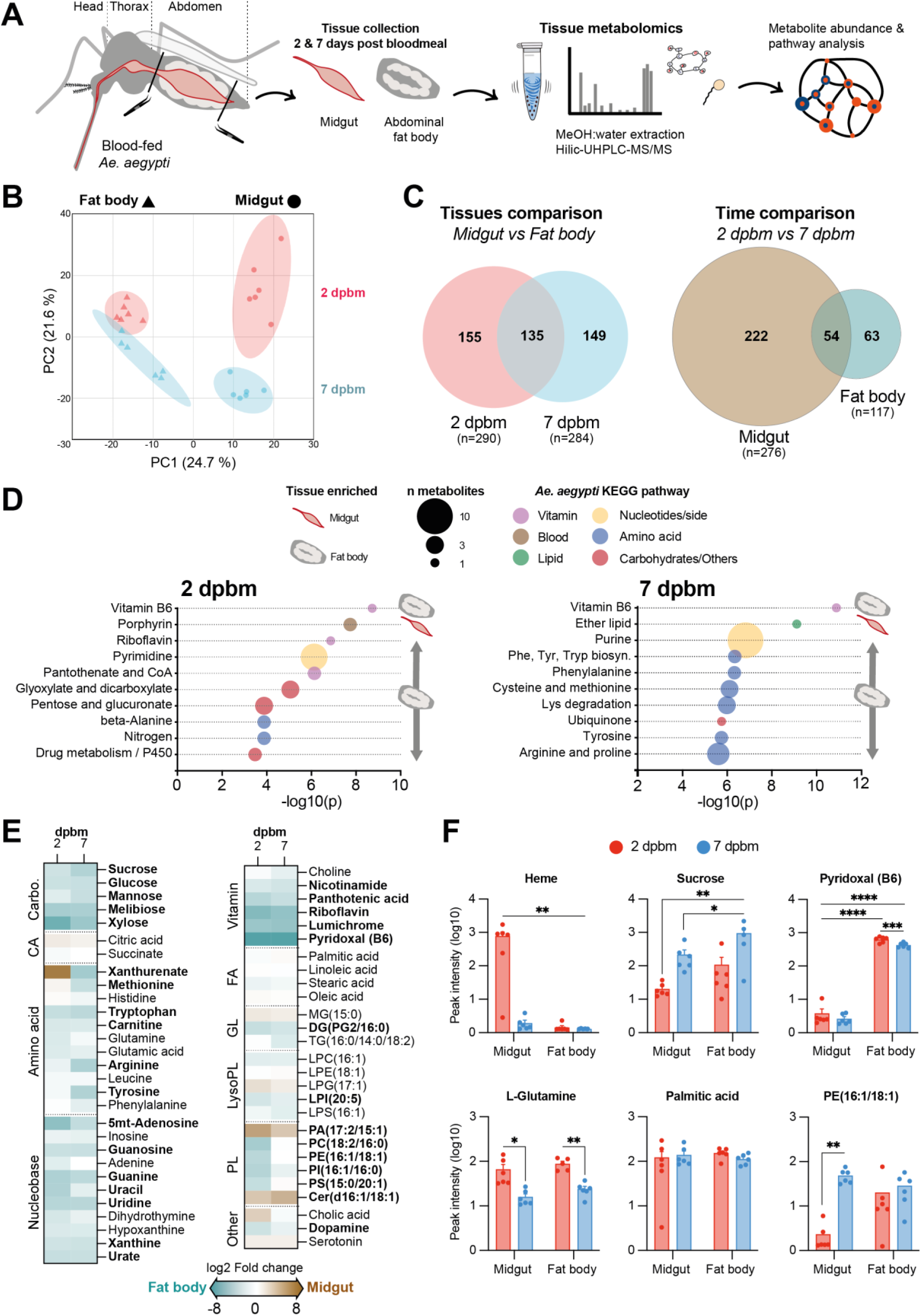
Tissue metabolomics highlight the physiological differences and the metabolic expertise of the fat body. (**A**) Graphical overview of the midgut and fat body collection on blood-fed mosquitoes for subsequent metabolites extraction using methanol: water (80:20) and metabolomics by UHPLC-MS/MS. Mosquitoes used for both metabolomic analysis and scRNAseq were from the same batch and experiments were conducted concurrently within the same facility. (**B**) Principal Component Analysis (PCA) of metabolomic dataset from dissected midgut and fat body tissues (**Supplementary file 6**). Each dot represents a pool of 5 tissues and pairs of midgut and fat body originated from the same individual. Samples were normalized by the total ion chromatogram. The dataset was log_10_ transformed and scaled by autoscaling. (**C**) Venn diagram of significantly regulated metabolites between midgut and fat body or between 2 and 7 days post-bloodmeal (dpbm). Significant metabolites were selected with a p-value FDR adjusted <0.05 and at least I2I fold change difference. (**D**) Pathway analysis of the 10 most different metabolic pathways between midgut and fat body at 2 and 7 dpbm. Pathway analysis used HMDB annotated metabolites and global test enrichment method, based on *Aedes aegypti* KEGG pathway library. Significant pathways were selected with -log_10_(p) >1,5 and p-value FDR adjusted <0.05. (**E**) Heatmap showing the differential abundance of selected metabolites between the midgut and the fat body at 2 and 7 dpbm. Significantly regulated metabolites in at least one time point are shown in bold and correspond to a fold change >I2I and p-value adjusted < 0.05, between midgut and fat body tissues. Metabolites are grouped by chemical classes. Carbo: carbohydrates; CA: carboxylic acids; FA: fatty acids; GL: glycerolipids; LysoPL: lysophospholipids, PL: phospholipids. (**F**) Selected metabolite intensity repartition in midgut and fat body at 2 and 7 dpbm. Significant differences were based on a two-way ANOVA Tukey’s multiple comparison p-value <0.05.

Pathway analysis reveals divergent metabolic pathway enrichment between tissues across time (**Fig. 5D**). The fat body exhibits consistent enrichment in vitamins, including sustained abundance in vitamin B6 (pyridoxal) and early abundance of riboflavin and pantothenate. This enrichment highlights a potential fat body role, notably for FYC (**Fig. 4**), in storing vitamins for subsequent utilization in metabolic processes as enzyme cofactors. Additionally, a gene associated with vitamin B6 metabolism (*Pdxk*), shows expression across tissues, including notable activity in EC4 and cardia cells within the midgut, as well as TP2 and HC-O in the fat body. This pattern highlights the fat body’s central role in supplying essential vitamins to dependent tissues and cells. Nucleotide metabolites are enriched in the fat body with an abundance shift observed from pyrimidine metabolism at 2 dpbm to purine metabolism at 7 dpbm, accompanied by a significant increase in amino acid metabolism. While systemic *awd* gene expression potentially supports nucleotide synthesis in the fat body at both time points, the high levels of amino acid metabolites observed at 7 dpbm in this tissue may result from increased expression in amino acid pathway genes in the midgut (**Fig.4**). In contrast, the midgut shows enrichment in porphyrin metabolism early on, followed by ether lipid metabolism at 7 dpbm.

We then identify notable differences in metabolite content between tissues, with the fat body displaying a higher abundance across most classes compared to the midgut (**Fig. 5E**). However, xanthurenate, a metabolite derived from the tryptophan/kynurenine pathway and known for its antioxidant and heme-chelating properties in mosquito midgut after a bloodmeal^88^, is highly abundant in the midgut at 2 dpbm and subsequently shifted to the fat body by 7 dpbm. In addition, the presence of heme exclusively in the midgut at 2 dpbm aligns with the breakdown of blood cells during the early stages of digestion (**Fig. 5F**). The fat body’s role in vitamin storage is further underscored by the high levels of vitamin detected, with B6 being particularly abundant (**Fig. 5E-F**).

Additionally, carbohydrate-related compounds, notably sucrose, are enriched in the fat body (**Fig. 5E-F**). This finding aligns with the expression of trehalose biosynthesis genes in fat body clusters (**Fig. 4**), given that trehalose synthesis requires sucrose as a substrate. Amino acids and nucleobase metabolites are slightly more abundant in the fat body (**Fig. 5E**). For instance, L-Glutamine, pivotal in metabolic and neuronal pathways, illustrates similar abundance in both tissues (**Fig. 5F**). This observation confirms that amino acid metabolism is as actively engaged in the midgut as it is in the fat body (**Fig. 4**). Levels of carboxylic acid and fatty acid show no significant differences in abundance between the midgut and fat body (**Fig. 5E**), suggesting that both tissues maintain comparable levels of these metabolites. Palmitic acid, a prevalent saturated fatty acid, is present at similar levels across tissues (**Fig. 5F**), despite fatty acid synthases being predominantly expressed in the fat body (**Fig. 4**), pointing to lipid exchanges between compartments. As for membrane lipids, the midgut is enriched in ceramide and phosphatidic acid (PA), a precursor for multiple lipid classes. (**Fig. 5E**). Notably, phospholipids with specific head groups characteristic of cell membranes (PC, PE, PI, PS), are significantly less abundant at 2 dpbm in the midgut, exemplified by phospholipid PE(18:1/16:1), (**Fig. 5F**). It suggests that bloodmeal digestion may have a profound impact on phospholipid remodeling within the midgut, a process known to intensify when the blood contains dengue virus^91^.

This comprehensive metabolomic analysis validates the distinct physiological states and specialized metabolic functions observed by scRNAseq on the midgut and the fat body of blood-fed *Ae. aegypti,* illustrating the specific metabolic pathways operational following a bloodmeal.

### Expression of insect-specific viruses

*Ae. aegypti* mosquitoes host a variety of microbial and viral communities, including several insect-specific viruses (ISVs)^92^. In our study, we aimed to detect the presence of well-described ISVs such as Phasi Charoen-like virus (PCLV), Humaita-Tubiacanga virus (HTV), and Cell-fusing agent virus (CFAV) in midgut and fat body cells. We quantitatively assessed the percentage of cells harboring viral RNA (**Fig. 6A**).

**Figure 6.**
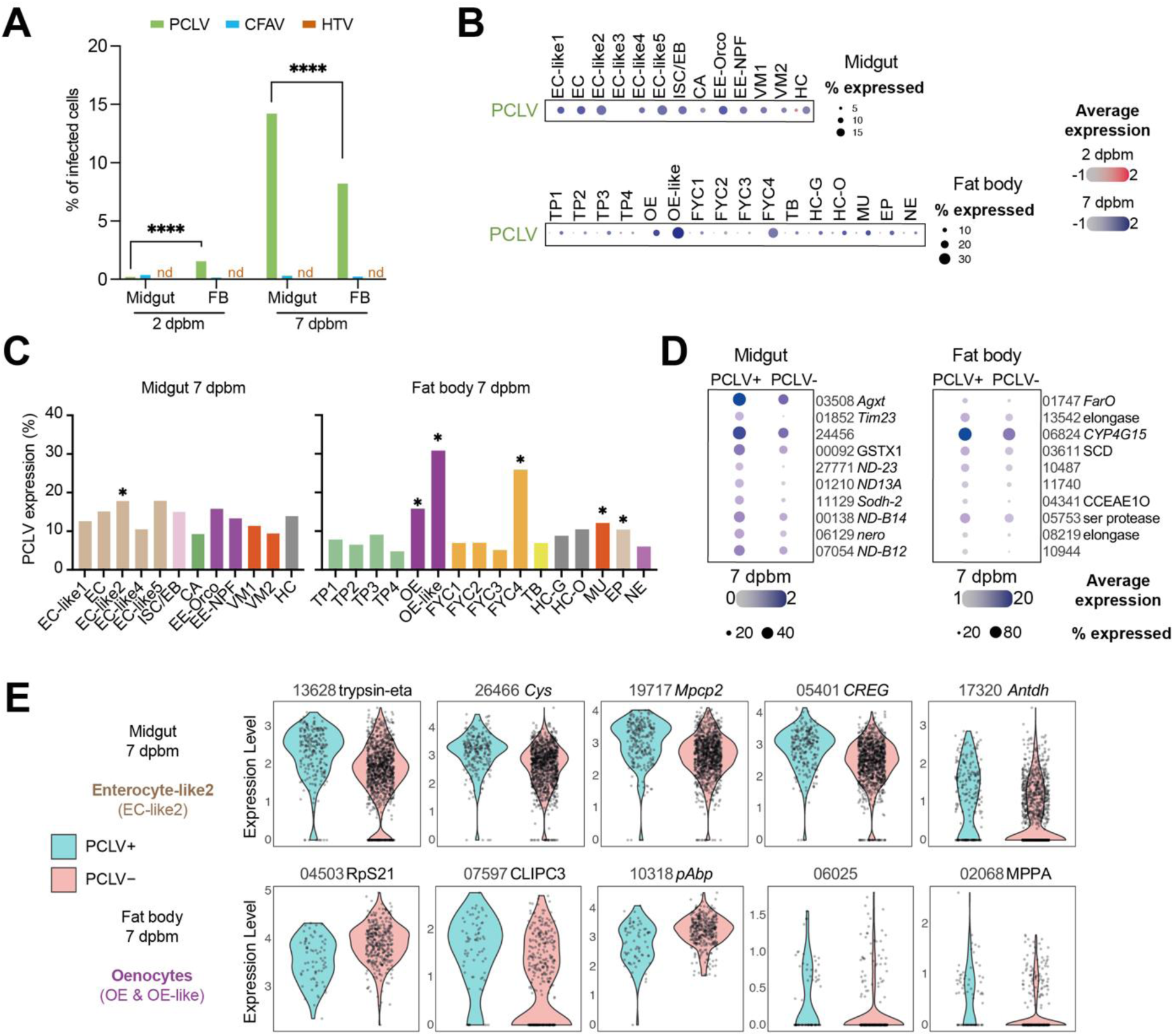
Insect-specific virus PCLV is highly expressed in midgut and fat body cells at later stage post-blood meal. (**A**) Proportion of cells expressing insect-specific viruses PCLV, CFAV and HTV in the midgut and the fat body, at 2 and 7 dpbm. nd: not detected. (**B**) Dot plot showing ISV expression of PCLV for midgut (top) and fat body (bottom) clusters. Dot color shows relative average expression intensity for ISVs at 2 dpbm (red) and 7 dpbm (blue). Dot size reflects the percentage of cells expressing PCLV in each cell cluster. (**C**) Proportion of cells expressing PCLV in each midgut and fat body clusters at 7 dpbm. Higher proportions of PCLV-positive cells were determined to be statistically significant by the chi-square test with Bonferroni correction, * = p<0.05. (**D**) Dot plot showing top 10 markers differentially expressed between PCLV-positive and PCLV-negative cells within midgut and fat body tissues. Dot color shows relative average gene expression intensity at 7 dpbm. Dot size reflects the percentage of cells expressing corresponding genes in each cell cluster. (**E**) Violin plot showing top 5 gene markers differentially expressed between PCLV-positive and PCLV-negative cells within enterocyte-like2 (EC-like2) and merged oenocytes (OE and OE-like) clusters, at 7 dpbm.

At 2 dpbm, PCLV transcripts are detectable in less than 2% of cells in both tissues. By 7 dpbm, however, 14% of midgut cells and 8% of fat body cells contain detectable levels of PCLV RNA, indicating a significant shift in infection, particularly in the midgut. Conversely, CFAV presence is consistently low, detected in less than 0.4% of cells across both time points and tissues, while HTV was absent in all samples (**Fig. 6B**). These data highlight insect-specific viral replication dynamics within mosquitoes, with an increase in PCLV viral replication following a bloodmeal, as previously reported in arbovirus-infected mosquitoes^93^.

Specifically, PCLV expression is slightly enriched in midgut EC-like2 cells and predominantly in fat body oenocytes (OE, OE-like) by 7 dpbm (**Fig. 6B-C**), and in a lesser extent in FYC4, MU and EP clusters. To explore how PCLV infection influences cellular function, we analyze gene expression differences between PCLV-positive and PCLV-negative cells at 7 dpbm (**Fig. 6D**). PCLV-infected midgut cells are enriched in genes associated with amino acid synthesis, oxidative respiration, and stress response (e.g., *serine-pyruvate aminotransferase*, *GST*, *NADH dehydrogenase*), indicative of EC-like2 activity. In the fat body, PCLV-positive cells upregulate lipid biosynthesis (*SCD*, *elongase*) and cytochrome P450 pathways (*CYP4G15*), which are characteristic of oenocytes. We then delve into the distinct transcriptional responses in the most infected cell cluster of each tissue (**Fig. 6E**). EC-like2 PCLV-positive cells display increased expression of genes associated in proteolysis modulation (*trypsin-eta, Cys*), ATP synthesis (*Mpcp2*), cellular homeostasis (*CREG*) and hormone synthesis (*Antdh*), supporting enhanced metabolic processes and energy production for viral replication. Meanwhile, PCLV-positive oenocytes (pooled Oe and Oe-like clusters), display a reduction in ribosomal protein expression (*RpS21*) and mRNA translation (*pAbp*) coupled with a mild upregulation of immune-associated serine proteases (*CLIPC3*) and mitochondrial metabolism (*MPPA*). These changes may constitute counter-defense mechanisms against viral invasion.

These results highlight the complex and specific responses of mosquito cells to endogenous virus infection, with a particular focus on PCLV in the midgut at 7 dpbm. The presence of PCLV prompts highly infected cells to display subtle transcriptional changes that indicate an active response to infection.

## DISCUSSION

Leveraging the power of single-cell transcriptomics and untargeted metabolomics led us to comprehensively characterize the tissular and cellular responses of the midgut and abdominal fat body of *Ae. aegypti* mosquitoes to a bloodmeal. Our cell atlases, including the first fat body cell atlas in arthropods, are available for exploration and analysis on an interactive online platform (https://aedes-cell-atlas.pasteur.cloud).

Our study reports 14,447 transcriptomes of individual mosquito midgut cells. We identify new marker genes for several cell populations, including enterocytes, enteroendocrine cells, intestinal stem cells/enteroblasts, cardia, visceral muscle cells, and associated hemocytes. While previous *Ae. aegypti* mosquito single-cell midgut studies focused on the effects of food (blood or sugar) or Zika virus infection, our analysis reveals the temporal transcriptional shifts in midgut cell populations at 2 and 7 days after the bloodmeal intake and digestion^15–17^. We performed a comparative analysis (**Fig. S2**) of all published data revealing an overall consistency in midgut cell type proportions and markers with other *Aedes* cell atlases and the reference *Drosophila* gut atlas^15–17,24^. Importantly, both single-nucleus RNA-Seq and scRNA-Seq yielded highly concordant results on midgut cells, suggesting that these approaches may be considered equivalent for future investigations^15^. Furthermore, marker genes identified in our study to associate with midgut cell clusters such as enterocytes (ECs), enteroendocrine cells (EE) and cardia cells (CA) aligned well with previously described bulk transcriptomic signatures of *Ae. aegypti* gut regions^30^. Together, the range of physiological states and the use of different *Ae. aegypti* strains (laboratory and field-collected) enrich our understanding of mosquito midgut cell composition while reinforcing the consistency of marker genes, cell types, and their proportions. However, accurately comparing cell numbers across conditions in single-cell studies is challenging, highlighting the need for improved validation methods in differential abundance analysis^94^.

Our study also provides the first fat body cell atlas published for arthropods, encompassing 27,726 individual transcriptomes. Our analysis uncovers a surprising level of cellular heterogeneity within the fat body, identifying well-known populations such as trophocytes and oenocytes, alongside neuroendocrine cells, muscle cells, tracheoblast-like cells, and epidermal cells^18,42,74,95–98^. We confirm the anticipated roles of trophocytes and oenocytes in vitellogenesis and metabolism, with trophocytes involved in hemolymph sugar synthesis and oenocytes in lipid biosynthesis. Additionally, our results support the proposed role of trophocytes in modulating antimicrobial humoral immunity, a critical fat body function observed as early as two days after the bloodmeal, likely influenced by the gut microbiome^43,46,99^. These observations also illustrate the tight links between metabolism and immunity, and their interplay with blood digestion mechanism.

Importantly, our analysis reveals previously unknown cell population in the fat body, that we called fat body-yolk cells. The gene expression profiles of fat body-yolk cells were reminiscent with those of ovarian cells^30,60,100,101^, that were however specifically removed during organ dissection. The significant and consistent abundance of fat body-yolk cells (18% of all cells at 2 and 7 dpbm) also discredits the hypothesis that fat body-yolk cells are simply residual ovarian cells. Rather, it suggests that fat body-yolk cells are inherent components of the abdominal fat body and play a critical role, possibly in close association with ovarian functions, in egg development. Specific fat body-yolk cells subpopulations (FYC2-3-4) express stem cell markers and showed enrichment of piRNA genes, similar to a germarium-like profile, which is essential for initiating egg chamber development^101,102^. Given the lack of oviposition sites during our experiment and the consistent presence of these cells over time, fat body-yolk cells might be instrumental in preserving egg integrity within mosquito females carrying mature eggs^69^. This finding challenges our current view of the fat body’s role in mosquito fertility.

The discovery and annotation of dozens of marker genes for mosquito cell population opens possibility for genetic tool development, such a tissue- and cell-specific drivers. Widely available in the model organism *D. melanogaster*, specific drivers combined with expression of fluorophores, apoptotic genes, or short-hairpin RNAs enable in-depth functional studies and elucidation of fundamental biological processes^103^. In mosquitoes, they might also lead to the development of the next generation of vector control tools, targeting only subsets of cells that are critical to pathogen transmission, and enable pathogen blockage with exquisite specificity, at each key step of the mosquito infectious cycle.

Our study constitutes a foundation for short- to long-term future investigations, aiming to validate the presence of distinct cell populations, such as fat body-yolk cells, using RNA *in situ* hybridization techniques or spatial transcriptomics. Thus, cell populations and states could be placed in the context of tissular architecture and a wider range of cellular interactions^104,105^. Such work could also provide a comprehensive view of the three-dimensional organization of mosquito abdominal tissues, shedding light on potential inter-organ interactions and metabolic gradients^106,107^. Recent advances in single-cell metabolomics also hold great promise for complementing single-cell transcriptomics data sets trough integrated multi-omics approaches^108^.

Hitherto, only one study has described the impact of an arbovirus infection, namely Zika virus, on mosquitoes at the single-cell level, and only on midguts 4 days post virus exposure. Building a comprehensive dataset, recapitulating the trajectory of a virus through a mosquito’s body, organ by organ, at the single-cell level would provide critical insights into the cellular mechanisms of pathogen susceptibility and transmission^17^. It could also shed light on the differential susceptibility and vectorial capacity of certain mosquito strains over others, informing spread of arboviruses and epidemiology. Moreover, in the context of the release of lab-modified mosquitoes carrying *Wolbachia*, an intracellular bacteria with virus-blocking properties in several regions of the globe, elucidating the fundamental processes underlying virus blockage at the single-cell level could provide insights into the heterogeneous efficiency rates observed on the field^109^.

Finally, enhancing our understanding of mosquito physiology at the cellular level, particularly in the fat body, is important for unraveling the mechanisms of insecticide resistance. Our study reveals that *Ae. aegypti* oenocytes robustly express key cytochrome P450 genes, essential for metabolic resistance to insecticides, aligning with observations in *Anopheles* and *Drosophila* that underscore the universal detoxification role of these cells^110^. Elucidating tissue-specific expression pattern of key insecticide resistance gene could lead to better identification and characterization of insecticide-resistant mosquito strains and development of counter strategies. Also, intriguing links between insecticide resistance and vector competence could be explored at the cellular and molecular level^111^.

In sum, the study of arthropod vectors and their interactions with symbionts, microbiomes and human pathogens at a single-cell resolution will hopefully generate deeper fundamental understanding of those complex interactions and pave the way towards the development of innovative disease control strategies.

## METHODS

### Mosquito rearing

Experiments were conducted with *Ae. aegypti* mosquitoes derived from a wild-type colony originating from Kumasi, Ghana, predominantly comprised of *Aedes aegypti formosus*, with a minor component of admixed ancestry from *Aedes aegypti aegypti*^19,20^. Mosquitoes were reared under controlled conditions (28°C, 12-hour light/12-hour dark cycle and 70% relative humidity). The mosquito colony was maintained by feeding female mosquitoes on rabbit blood (BCL), to ensure their survival, laying and egg collection. Prior to performing the experiments, their eggs were hatched synchronously in a SpeedVac vacuum device (Thermo Fisher Scientific) for 45 minutes. Their larvae were reared in plastic trays containing 1.5 L of tap water and supplemented with Tetramin (Tetra) fish food at a density of 200 larvae per tray. After emergence, adults were kept in BugDorm-1 insect cages (BugDorm) with permanent access to 10% sucrose solution.

### Mosquito bloodmeal

Five- to 7-day-old female mosquitoes were deprived of 10% sucrose solution 18-24 hours before exposure to an artificial bloodmeal consisting of a 2:1 mix of washed rabbit erythrocytes (BCL) supplemented with 10 mM adenosine triphosphate (Sigma) and Leibovitz’s L-15 medium (Gibco) supplemented with supplemented with 0.1% penicillin/streptomycin (pen/strep; Gibco ThermoFisher Scientific), 2% tryptose phosphate broth (TBP; Gibco Thermo Fischer Scientific), 1× non-essential amino acids (NEAA; Life Technologies) and 10% fetal bovine serum (FBS; Life Technologies). Mosquitoes were exposed to the bloodmeal for 15 min through a desalted pig-intestine membrane using an artificial feeder (Hemotek Ltd) set at 37°C. Fully engorged females were placed into small containers and maintained for 7 days at 28°C, 70% relative humidity and under a 12-hour light-dark cycle with permanent access to 10% sucrose. To meet biosafety requirements, containers did not provide space for oviposition, effectively preventing egg-laying.

### Dissection of midgut and fat body

Midgut dissection consisted of isolating the digestive track from the mosquito’s body by gripping the second to last abdominal segment using a forceps and pulling it out. Only the midgut, devoid of the crop, foregut, hindgut and Malpighian tubules was kept for analysis. Fat body dissection consisted of collecting the dorsal and ventral abdomen to which the fat body lobes are attached. Crop, Malpighian tubules, hindgut and ovaries were removed if needed.

### Single-cell dissociation of midgut and fat body

A cold dissociation method was used for the mosquito midgut and fat body, adapting protocols originally developed for dissociating *Anopheles sp*. midgut into single-cell suspensions suitable for scRNAseq^22^. Mosquitoes were dissected on ice and pools of 7 organs collected in a drop of ice-cold SF900 III media (Sigma-Aldrich). Each organ was sectioned into 3 smaller pieces using spring scissors (Fine science tools). Sectioned organs were transferred immediately to a chambered coverglass well (Thermo scientific) containing 500ul of dissociation media (SF900 III media supplemented with 10mg/ml Bacillus licheniformis protease (Sigma-Aldrich) and 25 unit/ml DNaseI (Sigma)) for 15 minutes in the dark on ice. Tissues were then gently triturated using 15 back-and-forth pipetting motions. After 5 minutes, 100uL of supernatant containing dissociated cells was collected and transferred to a collection tube containing 45 mL of SF900 III media on ice. The dissociation well was supplemented with 100uL of fresh dissociation media and incubated for 10 min in the dark on ice. Tissues were triturated again, and 100uL of supernatant transferred to the collection tube. Three to four more dissociation rounds were performed until only single cells remained. The collection tube was then spun at 300g for 15 min at 4°C. The pellet was resuspended gently in 900 ul ice-cold SF900 II media and filtered through Flowmi cell strainers (Sigma) with a 40 um porosity for midguts and 70 um porosity for fat bodies. The resulting cell suspension was spun at 300 g for 5 min at 4°C and resuspended in 30 ul of PBS 1x-BSA 0,04%. Cell concentration was determined using a Kova slide, and final concentration adjusted to 700-1000 cells per microliter.

### Single-cell gene expression assay and sequencing

Single-cell RNA sequencing was performed using the Chromium Next GEM Single Cell 5’ Kit v2 dual index (10x Genomics) according to the manufacturer’s protocol. Following a protocol developed in the context of a pilot study, a primer specific to the dengue virus envelope sequence was added at a concentration of 1uM, (Sequence 5’-3’: AAGCAGTGGTATCAACGCAGAGTACGAACCTGTTGATTCAACAGC). The targeted recovery rate was 10,000 cells per sample. Sample quality control was performed at Omics core facility at Institut Pasteur using Bioanalyzer DNA1000 high sensitivity (Agilent) and Qubit 4 Fluorometer (Thermo Fisher Scientific) assays. Libraries were sequenced by Macrogen on a Novaseq 6000 platform (Illumina).

### Reference generation and sample processing with STARsolo

As a reference genome for *Aedes aegypti*, we use the AaegL5 genome from Vectorbase (NCBI GenBank assembly id: GCA_002204515.1 https://www.ncbi.nlm.nih.gov/datasets/genome/GCF_002204515.2/). The fastas and the gtfs from *Aedes aegypti* were merged in one fasta file and one gtf respectively. The merged fasta and gtf were used by STAR to generate the index files with the command^112^. The index files were generated with the command STAR --runMode genomeGenerate using the gtf file as sjdbGTFfile and the fasta file as genomeFastaFiles. All sequencing data were processed and mapped to the Ae. aegypti with STAR (version 2.7.9a) using the following parameters: -- soloType CB_UMI_Simple --soloBarcodeMate 1 --clip5pNbases 42 0 --soloCBstart 1 --soloCBlen 16 --soloUMIstart 17 --soloUMIlen 10 --outFilterMultimapNmax 50 --outFilterScoreMinOverLread 0.4 --outFilterMatchNminOverLread 0.4 --outBAMcompression 10 --soloMultiMappers EM --soloStrand Forward.

### Single-cell quality control and cell filtering

STARsolo raw output files were individually read into RStudio (R version 4.3.2 from 2023-10- 31) and processed using Seurat (Seurat v5.0.1) and SingleCellExperiment (version 1.24.0). Samples were converted to Seurat objects and SingleCellExperiment objects. A first filter was applied by using the DropletUtils::barcodeRanks function (DroptletUtils version 1.22.0): the knee point of the barcoderank was used as a cutoff to remove empty and /or bad quality droplets with a total UMI number under this value (**Table S6**). After this first filter, we obtained for samples Midgut-2dpbm, Midgut-7dpbm, Fatbody-2dpbm and Fatbody-7dpbm respectively 33,832, 49,826, 46,158 and 48,205 cells that is at least twice as much as the number of cells targeted (20000 cells). Samples were converted to Seurat objects. Mitochondrial gene percentage was calculated for each cell. Features were log normalized, variable features were identified, the top 2,000 variable features were scaled, a principal component (PC) analysis dimensionality reduction was run. Nearest neighbors were computed, and appropriate clustering granularity was determined with Clustree (v0.5.1). Uniform Manifold Approximation and Projection (UMAP) dimensional reduction was performed, and clusters were visualized with UMAP reduction. In each sample, a second filter for cells was applied by removing cells contained in clusters 0 and 1, as these clusters have, in all samples, few to no significant marker genes and/or mitochondrial genes as genes markers (**Table S6**).

We also applied the following criteria to exclude poor quality cells and potential doublets or multiplets^15,24^, which can display aberrantly high gene counts and do not represent true single cells:

- Cells were retained if they had more than 100 detected genes.
- For midgut samples, cells with fewer than 2,400 detected genes were kept, based on the 99th percentile distribution.
- For fat body samples, cells with fewer than 5,300 detected genes were kept, following the same percentile-based exclusion.
- Mitochondrial genes were filtered with the following criteria: cells with more than 5% mitochondrial genes were excluded.

These metrics ensured a cell complexity greater than 0.75, defined as the log10(genes detected) divided by the log10(UMI counts per cell), and no cells retained for further analysis contained fewer than 400 total reads.

### Seurat workflow

Raw counts for cells passing all our filters were loaded into a Seurat object. For midgut and fat body, both samples at day 2 and day 7 were merged (Seurat::merge). Samples from the same tissue were integrated, following the integration tutorial proposed by Seurat authors (https://satijalab.org/seurat/articles/seurat5_integration). Features were log normalized, the top 2000 variable features were identified. All variable features were scaled and a principal component (PC) analysis dimensionality reduction was run. We integrated both samples (day 2 and day 7) together for midgut and fat body tissues using the CCAIntegration method from Seurat. After integration, nearest neighbors were computed, and appropriate clustering granularity was determined with Clustree (res 0.5 and 0.3 for midgut and fat body respectively). Uniform Manifold Approximation and Projection (UMAP) dimensional reduction was performed (parameters: n.neighbors=100, min.dist = 0.3), and clusters were visualized with UMAP reduction.

### Metabolite extraction

Mosquitoes were dissected in PBS 1X, and tissues collected in pools of 5 in tubes containing 500 ul of ice-cold methanol:water (80:20) and ∼20 1-mm glass beads (BioSpec). Samples were homogenized for 30 sec at 6,000 revolutions per minute (rpm) in a Precellys 24 grinder (Bertin Technologies), and sonicated in an ultrasonic bath (J.R. Selecta) for 15 min at 4 °C. Homogenates were centrifuged at 10,000 rpm for 1 min at 4 °C and 400 µL of supernatant collected. Remaining pellets were resuspended in 500 µL of methanol:water (80:20) solution. After another round of sonication and centrifugation, 400 µL of supernatant was collected. Combined supernatants were vacuum-dried using Speed-Vac for 4 hours (Thermo Fisher Scientific) and stored at −20 °C. Four to six biological replicates consisting of pools of 5 tissues were prepared per condition.

### Liquid Chromatography-High Resolution Mass Spectrometry (LC-HRMS) Metabolic Profiling

Dry extracts were resuspended at 2 mg/mL in an 80:20 methanol:water solution. Ultra-high-performance liquid chromatography−high-resolution mass spectrometry (UHPLC−HRMS) analyses were performed on a Q Exactive Plus quadrupole (Orbitrap) mass spectrometer, equipped with a heated electrospray probe (HESI II) coupled to a U-HPLC Vanquish H (Thermo Fisher Scientific, Hemel Hempstead, U.K.). Hilic separation mode was done using a Poroshell 120 Hilic-Z (2.7µm, 150 × 2.1 mm, Agilent, Santa Clara, USA) equipped with a guard column. The mobile phase A (MPA) was water with 0.1% formic acid (FA) and 10 mM ammonium formate, and the mobile phase B (MPB) was acetonitrile with 0.1% FA and 10 mM ammonium formate. The solvent gradient was 10% MPA (0 – 0.5 min), 10% MPA to 50% MPA (0.5 - 9 min), 50% MPA (9 - 11 min), 10% MPA (11.1 - 20 min). The flow rate was 0.3 mL/min, the column temperature was set to 40 °C, autosampler temperature was set to 5 °C, and injection volume fixed to 1 μL. Mass detection was performed in positive and negative ionization (PI and NI) mode at resolution 35 000 power [full width at half-maximum (fwhm) at 400 m/z] for MS1 and 17 500 for MS2 with an automatic gain control (AGC) target of 1 × 10^6^ for full scan MS1 and 1 × 10^5^ for MS2. Ionization spray voltages were set to 3.5 kV for PI and 2,6 kV for NI, and the capillary temperature was kept at 256 °C. The mass scanning range was m/z 100−1500. Each full MS scan was followed by data-dependent acquisition of MS/MS spectra for the four most intense ions using stepped normalized collision energy of 20, 40, and 60 eV.

### Metabolomic data processing

UHPLC-HRMS raw data were processed with MS-DIAL version 5.1^113^ for mass signal extraction between 100 and 1500 Da from 0.5 to 12 min. Respectively MS1 and MS2 tolerance were set to 0.01 and 0.05 Da in centroid mode. The optimized detection threshold was set to 1 × 105 concerning MS1 and 10 for MS2. Peaks were aligned on a quality control (QC) reference file with a retention time tolerance of 0.1 min and a mass tolerance of 0.015 Da. All MS-CleanR filters were applied with RSD tolerance of 50% and blank suppression ratio of 0.8^114^. Peak annotation was performed with in-house database (level 1) and FragHub database (level 2) and finally using from MS-FINDER internal DB (level 3)^115,116^.

### Insect-specific virus detection

To get the ISV count, we performed a second run of read alignment using STAR (solo mode), counting reads on both strands (--soloStrand Unstranded) on a reference genome containing:

- AaegL5 genome (NCBI GenBank assembly id: GCA_002204515.1 - https://www.ncbi.nlm.nih.gov/datasets/genome/GCF_002204515.2/)
- CFAV (Cell fusing agent virus) strain Galveston, complete genome (https://www.ncbi.nlm.nih.gov/datasets/genome/GCF_000862225.3/ GenBank assembly id : GCA_000862225.1)
- HTV (Humaita-Tubiacanga virus) strain Ab-AAF complete genome (two segments) (https://www.ncbi.nlm.nih.gov/datasets/genome/GCA_031535205.1/ - GenBank assembly id GCA_031535205.1)
- PCLV (Phasi Charoen-like virus) isolate Rio, complete genome (three segments) (https://www.ncbi.nlm.nih.gov/datasets/genome/GCF_002814835.1/ - GenBank assembly id : GCA_002814835.1)

ISV counts were added as metadata for fat body and midgut cells. These counts were not considered for normalization, scaling, dimensional reduction or clustering steps.

### Statistical analysis

Statistical analyses were performed in GraphPad Prism version 10.2.2. Difference in cell clusters proportions and ISV-infected cell proportions were measured by chi-squared test. Difference in metabolite intensity were measured by 2-way ANOVA and multiple comparison test. Metabolomic dataset was analyzed using MetaboAnalyst 6.0 (www.metaboanalyst.org)^117^ and data were normalized by log transformation and auto scaling, for principal component analysis (PCA). Venn diagram were produced using InteractiVenn^118^, based on significant metabolites (fold changes >2, DFR adjusted p-value <0,05). Pathway analysis was based on well-annotated HMDB compounds for global test enrichment on *Aedes aegypti* (yellow fever mosquito) (KEGG).

## Data availability

The scRNAseq data discussed in this publication have been deposited in NCBI’s Gene Expression Omnibus and are accessible through GEO Series accession number available upon request. The metabolomic data discussed in this publication have been deposited in Zenodo and are accessible through the DOI 10.5281/zenodo.13990385.

## ACKNOWLEDGMENTS

We thank Louis Lambrechts for his supportive role and insightful guidance during the study, Sampson Otoo for helping establishing mosquito colony, Catherine Lallemand for assistance with mosquito rearing, Cassandra Koh for discussions about insect-specific viruses, Bryan Brancotte, Juliette Bonche and Rachel Torchet for website development, the Single Cell Biomarkers UTechS platform (C2RT, Institut Pasteur, Paris, France), in particular Milena Hasan, Valentina Libri, Carolina Moares-Cabe and Valérie Seffer for support on 10X Chromium and library preparation. The scRNA-seq library quality control and preparation for sequencing was performed by Elodie Turc from the Biomics platform (C2RT, Institut Pasteur, Paris, France) supported by France Génomique (ANR-10-INBS-09) by the French National Facility in Metabolomics & Fluxomics, MetaboHUB (11-INBS-0010) and IBISA. This work was supported by the French Government’s Investissement d’Avenir program, Laboratoire d’Excellence Integrative Biology of Emerging Infectious Diseases (ANR-LBX-IBEID S2I grant to S.H.M.), Agence Nationale de la Recherche (ANR JC/JC S-CR21073 grant to S.H.M.), L’Oréal For Women in Science Foundation (to S.H.M), Institut Pasteur (Post-doctoral Pasteur Roux-Cantarini fellowship to T.V.) and a European Union Horizon 2020 Marie Sklodowska-Curie Action (postdoctoral fellowship to T.V.). The funders had no role in study design, data collection and analysis, decision to publish, or preparation of the manuscript.

## AUTHOR CONTRIBUTIONS

Conceptualization: T.V., H.L.M., E.C., S.P., V.H., J.A., M.L., G.M. S.H.M.

Investigation: T.V., H.L.M., E.C, S.H.M.

Data curation: T.V, H.L.M.

Formal analysis: T.V., H.L.M., S.H.M.

Visualization: T.V., H.L.M., S.H.M.

Writing – original draft: T.V., H.L.M., S.H.M.

Writing – review and editing: T.V., H.L.M., E.C., S.P., J.A., M.L., G.M. S.H.M.

Funding acquisition: T.V., S.H.M.

## SUPPLEMENTARY FIGURES AND LEGENDS

**Figure S1.**
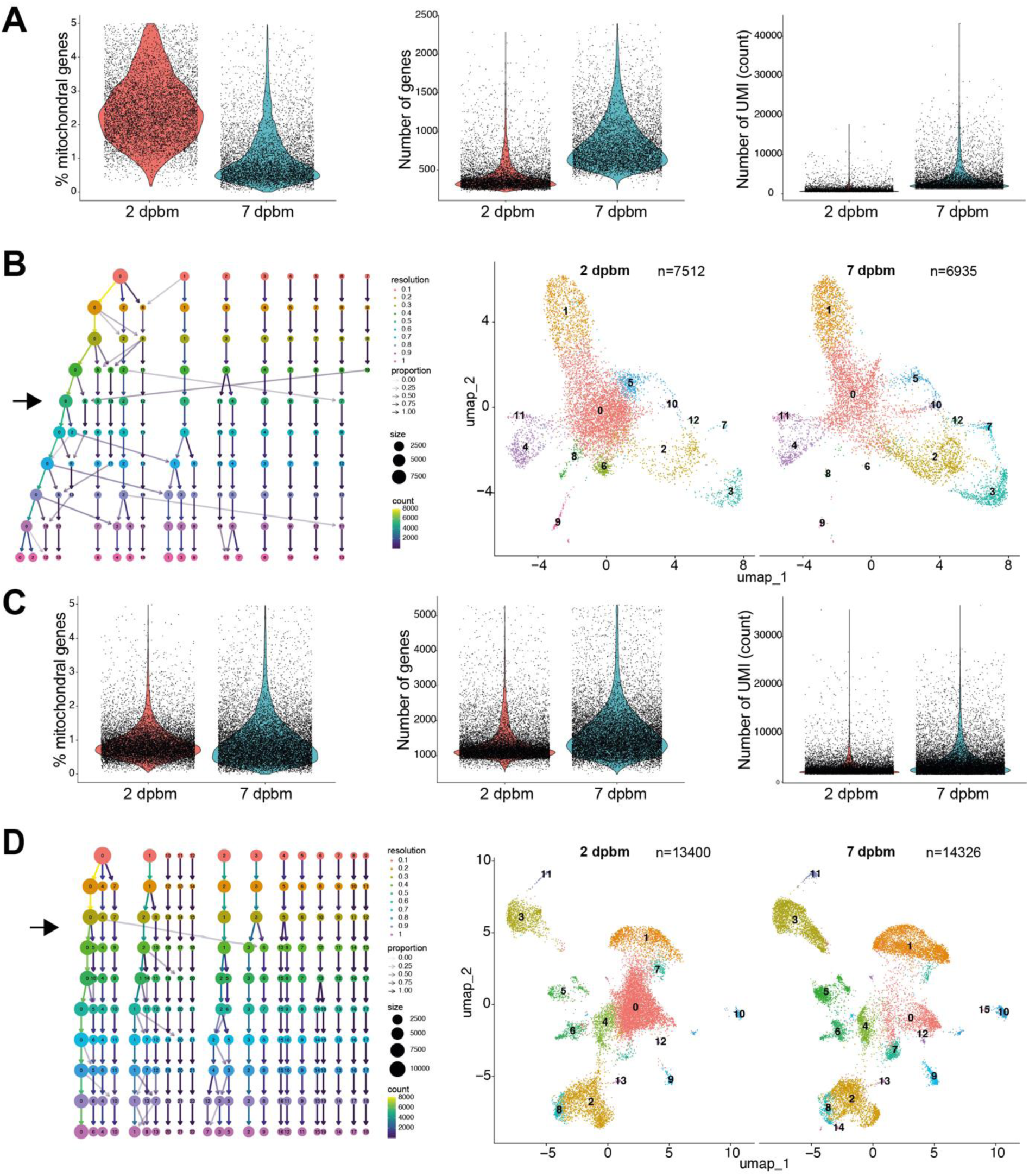
Quality control and clustering analysis of midgut and fat body cell samples. (**A**) Quality control metrics for midgut cells with mitochondrial gene proportion, gene count, and UMI count per cell at 2 and 7 dpbm. (**B**) Clustree visualization and UMAP analysis of midgut cell clustering across integrated 2 and 7 dpbm time-points, with chosen Seurat resolution of 0.5. (**C**) Quality control metrics for fat body cells with mitochondrial gene proportion, gene count, and UMI count per cell at 2 and 7 dpbm. (**D**) Clustree visualization and UMAP analysis of fat body cell clustering across integrated 2 and 7 dpbm time-points, with chosen Seurat resolution of 0.3.

**Figure S2.**
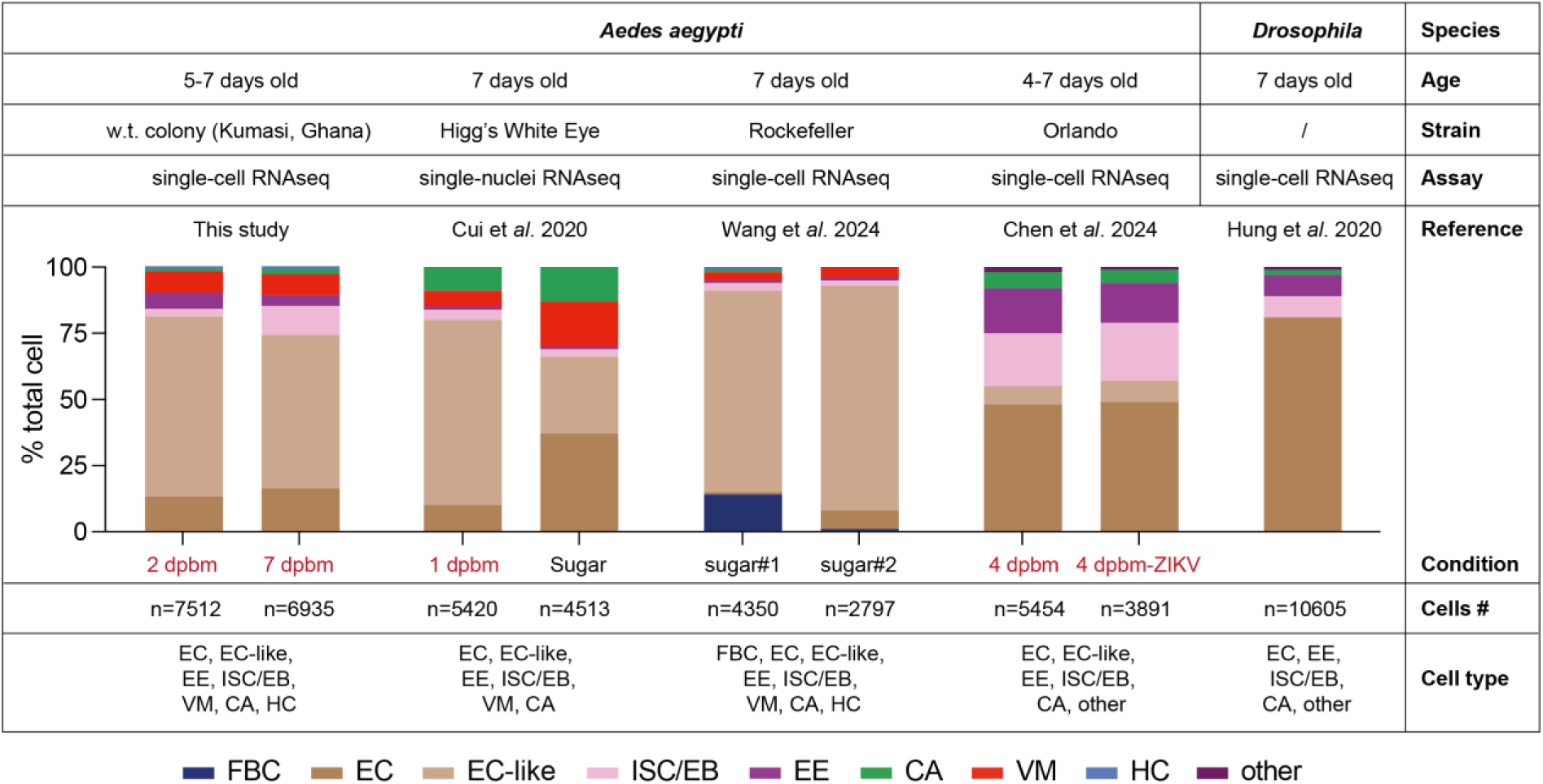
Comparison of female *Ae. aegypti* midgut cell composition with reported *Ae. aegypti* and *Drosophila* midgut cell studies. FBC: fat body cells; EC: enterocytes; ISC/EB: intestinal stem cells/enteroblasts; EE: enteroendocrine cells; CA: cardia cells; VM: visceral muscle cells; HC: hemocytes; w.t.: wild-type; dpbm: day post-blood meal.

**Figure S3.**
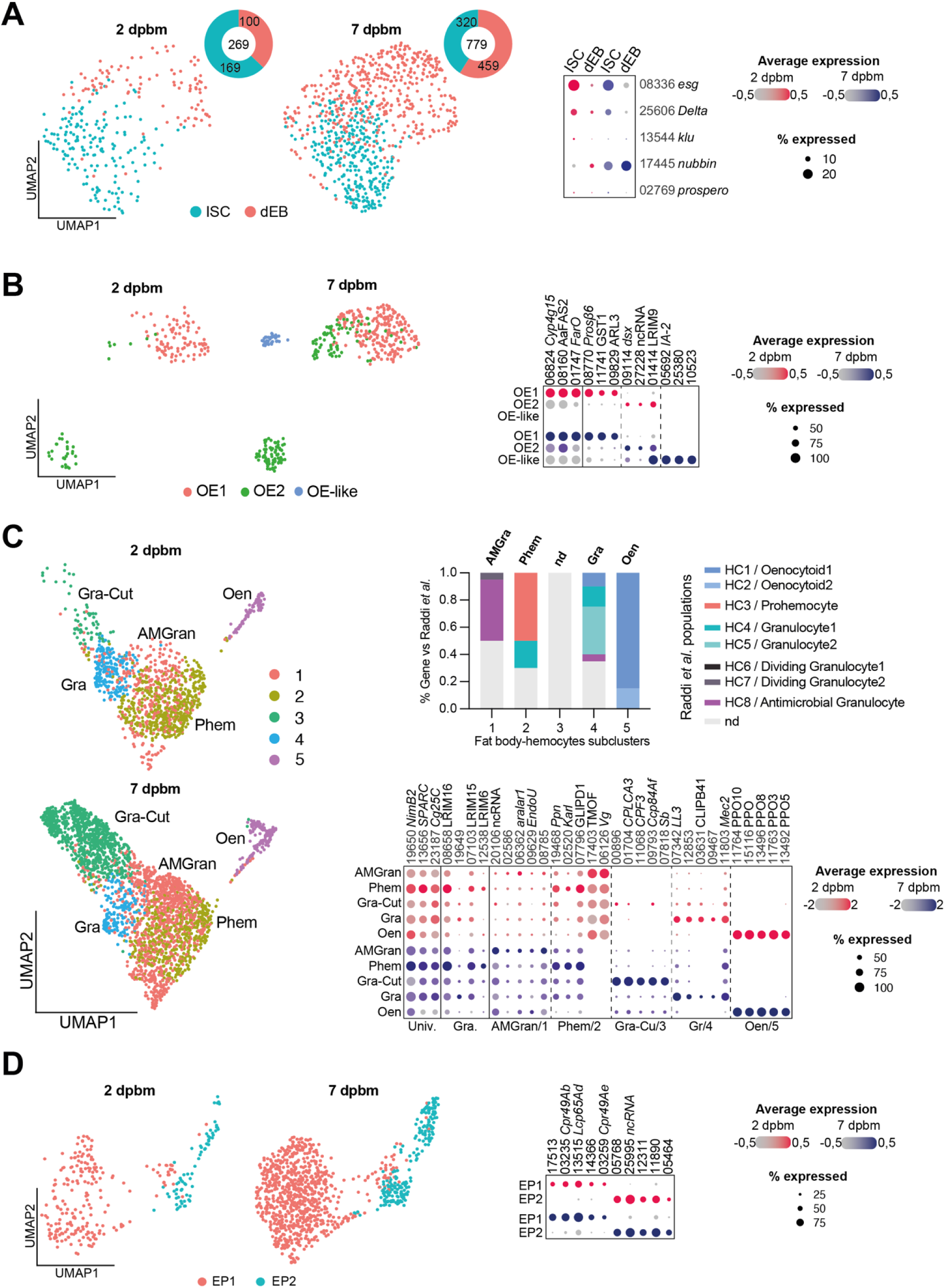
Sub-clustering improves characterization of differentiated cells within midgut and fat body populations. (**A**) UMAP visualization of intestinal stem cell/enteroblast (ISC/EB) group-specific clustering within the midgut at 2 and 7 dpbm using 0.2 resolution, separating ISC and differentiated EB (dEB) with their proportions at both time point by part of whole plot. Canonical stem cell gene markers (*esg, delta, klu*) and markers for differentiating EB (*nubbin, prospero*) are displayed in a dot plot. (**B**) UMAP visualization of oenocyte (Oe and Oe-like) group-specific clustering within the fat body at 2 and 7 dpbm using 0.1 resolution. The dot plot displays gene expression for highly specific marker presented in Fig. 3D and the top 3 gene markers from each oenocyte sub-cluster. (**C**) UMAP visualization of hemocyte group-specific clustering within the fat body at 2 and 7 dpbm using 0.3 resolution. Clusters are annotated based on the expression of genes matching those reported in hemocyte subpopulations by Raddi *et al*^46^. The bar plot illustrates the proportion of gene expression within each sub-cluster that corresponds to expression profiles in Raddi’s hemocyte populations. The dot plot displays gene expression for universal (Univ.) and granulocyte (Gra.) markers, along with the top 5 gene markers from each hemocyte sub-cluster. (**D**) UMAP visualization of epidermal cell (EP) group-specific clustering within the fat body at 2 and 7 dpbm using 0.2 resolution. The dot plot displays gene expression of the top 5 gene markers from each epidermal cell sub-cluster. Dot color shows relative average expression intensity for each gene at 2 dpbm (red) and 7 dpbm (blue). Dot size reflects the percentage of cells expressing corresponding genes in each cell cluster. Each gene is identified by the AAEL0 truncated Vectorbase IDs and their respective abbreviation if available, not italicized. *Drosophila* or *Anopheles* ortholog abbreviations are indicated in italic.

**Figure S4.**
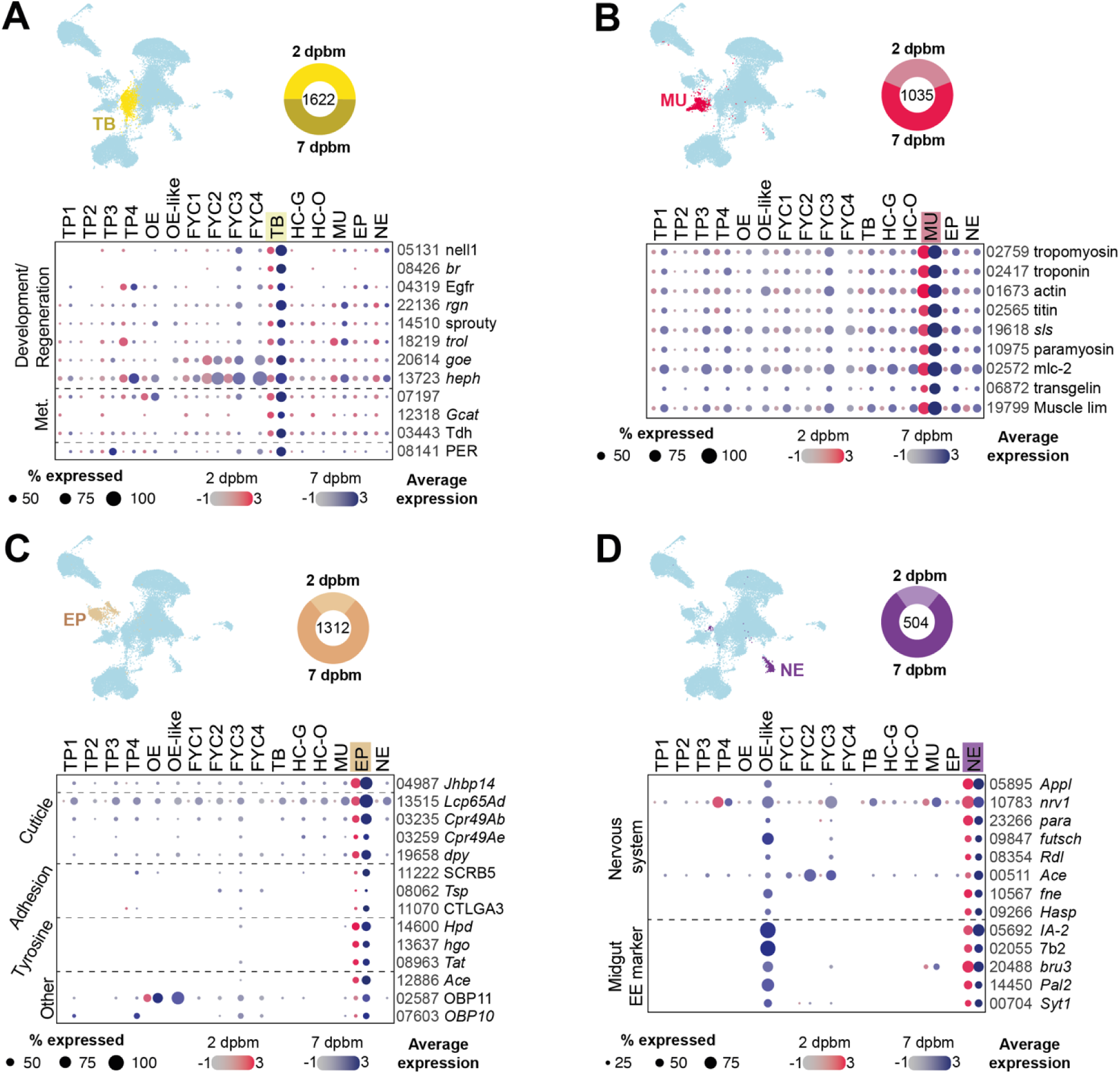
Heterogeneous and specialized cell populations in the fat body-associated tissues. Specific cell clusters visualized on UMAP associated with their cell proportions by part of whole plot and gene markers expression representative of (A) tracheoblast-like cells, (B) muscle cell (C), epidermal cell and (D) neuroendocrine cells. Dot color shows relative average expression intensity for each gene at 2 dpbm (red) and 7 dpbm (blue). Dot size reflects the percentage of cells expressing corresponding genes in each cell cluster. Each gene is identified by the AAEL0 truncated Vectorbase IDs and their respective abbreviation if available, not italicized. *Drosophila* or *Anopheles* ortholog abbreviations are indicated in italic. Met: metabolism; TB: tracheoblast; MU: muscle cells; EP: epidermal cells; NE: neuroendocrine cells.

## SUPPLEMENTARY TABLES

**Table S1.** Number of cells collected in midgut and fat body samples.

| Samples | Day of collection | N cells total | N genes detected >1 | genome coverage* | Median UMI/cell | Median Gene/cell |
| --- | --- | --- | --- | --- | --- | --- |
| **Midgut** | 2 | 7512 | 11358 | 57% | 759 | 360 |
|  | 7 | 6935 | 11690 | 59% | 2707 | 776 |
| **Fat body** | 2 | 13400 | 14170 | 72% | 2320 | 1184,5 |
|  | 7 | 14326 | 15162 | 77% | 3373,5 | 1506 |

\* (Aaeg L5.3: 14,718 protein coding genes, 4704 ncRNA genes, and 382 pseudogenes from Vectorbase)<sup>119</sup>

**Table S2.** Proportion of cells grouped into cell types in the midgut.

| Cell group | 2 dpbm | 7dpbm | 2 dpbm | 7dpbm | Chi-squared test |
| --- | --- | --- | --- | --- | --- |
| **EC-like** | 5106 | 4025 | 68,0% | 58,0% | 0.0008 |
| **EC** | 959 | 1123 | 12,8% | 16,2% | <0.0001 |
| **ISC/EB** | 269 | 779 | 3,6% | 11,2% | <0.0001 |
| **VM** | 568 | 543 | 7,6% | 7,8% | 0.95 |
| **EE** | 483 | 288 | 6,4% | 4,2% | <0.0001 |
| **CA** | 16 | 141 | 0,2% | 2,0% | 0.563 |
| **HC** | 111 | 36 | 1,5% | 0,5% | 0.763 |
| **Total** | 7512 | 6935 |  |  |  |

**Table S3.** Proportion of cells grouped into cell types in the fat body.

| Cell group | 2 dpbm | 7dpbm | 2 dpbm | 7dpbm | Chi-squared test |
| --- | --- | --- | --- | --- | --- |
| **TP** | 8007 | 5630 | 59,8% | 39,3% | 0.003 |
| **FYC** | 2470 | 2699 | 18,4% | 18,8% | 0.87 |
| **HC** | 1206 | 2707 | 9,0% | 18,9% | <0.0001 |
| **TB** | 814 | 808 | 6,1% | 5,6% | 0.59 |
| **EP** | 286 | 1026 | 2,1% | 7,2% | <0.0001 |
| **MU** | 392 | 643 | 2,9% | 4,5% | <0.0001 |
| **NE** | 103 | 401 | 0,8% | 2,8% | <0.0001 |
| **OE** | 122 | 412 | 0,9% | 2,9% | <0.0001 |
| **Total** | 13400 | 14326 |  |  |  |

**Table S4.** Proportion of midgut cells by cluster.

| Cell cluster | cluster | 2 dpbm | 7 dpbm | 2 dpbm | 7 dpbm |
| --- | --- | --- | --- | --- | --- |
| EC-like1 | 0 | 4112 | 2688 | 54,7% | 38,8% |
| EC | 1 | 959 | 1123 | 12,8% | 16,2% |
| EC-like2 | 2 | 558 | 1270 | 7,4% | 18,3% |
| ISC/EB | 3 | 269 | 779 | 3,6% | 11,2% |
| VM1 | 4 | 531 | 447 | 7,1% | 6,4% |
| EE-Orco | 5 | 447 | 190 | 6,0% | 2,7% |
| EC-like3 | 6 | 312 | 1 | 4,2% | 0,0% |
| CA | 7 | 16 | 141 | 0,2% | 2,0% |
| EC-like4 | 8 | 115 | 38 | 1,5% | 0,5% |
| HC | 9 | 111 | 36 | 1,5% | 0,5% |
| EE-NPF | 10 | 36 | 98 | 0,5% | 1,4% |
| VM2 | 11 | 37 | 96 | 0,5% | 1,4% |
| EC-like5 | 12 | 9 | 28 | 0,1% | 0,4% |
| total |  | 7512 | 6935 |  |  |

**Table S5.** Proportion of fat body cells by cluster.

| Cell cluster | cluster | 2 dpbm | 7 dpbm | 2 dpbm | 7 dpbm |
| --- | --- | --- | --- | --- | --- |
| TP1 | 0 | 6350 | 1737 | 47,4% | 12,1% |
| TP2 | 1 | 1354 | 3161 | 10,1% | 22,1% |
| FYC1 | 2 | 2058 | 2247 | 15,4% | 15,7% |
| HC-G | 3 | 1129 | 2554 | 8,4% | 17,8% |
| TB | 4 | 814 | 808 | 6,1% | 5,6% |
| EP | 5 | 286 | 1026 | 2,1% | 7,2% |
| MU | 6 | 392 | 643 | 2,9% | 4,5% |
| TP3 | 7 | 275 | 606 | 2,1% | 4,2% |
| FYC2 | 8 | 350 | 359 | 2,6% | 2,5% |
| NE | 9 | 103 | 401 | 0,8% | 2,8% |
| OE | 10 | 122 | 373 | 0,9% | 2,6% |
| HC-O | 11 | 77 | 153 | 0,6% | 1,1% |
| TP4 | 12 | 28 | 126 | 0,2% | 0,9% |
| FYC3 | 13 | 61 | 39 | 0,5% | 0,3% |
| FYC4 | 14 | 1 | 54 | 0,0% | 0,4% |
| OE-like | 15 | 0 | 39 | 0,0% | 0,3% |
| Total |  | 13400 | 14326 |  |  |

**Table S6.** Single-cell quality control process and cell filtering.

|  | Midgut-2dpbm | Midgut-7dpbm | Fatbody-2dpbm | Fatbody-7dpbm |
| --- | --- | --- | --- | --- |
| Inflection | 109 | 129 | 202 | 125 |
| knee | 484 | 1,248 | 1,597 | 1,528 |
| Cells after knee cutoff | 33,832 | 49,826 | 46,158 | 48,205 |
| Resolution for clustering (filtering step) | 0.4 | 0.35 | 0.3 | 0.3 |
| Nb cells after removing cluster 0+1 | 8,010 | 6,991 | 13,474 | 14,744 |
| Final cell number (filter percent.mt + min/max<br>number of genes) | 7,512 | 6,935 | 13,400 | 14,326 |

## SUPPLEMENTAL INFORMATION

Supplementary file 1: List of genes and orthologs used in this study.

Supplementary file 2: Significant markers of midgut cell clusters identified by FindAllMarkers.

Supplementary file 3: Significant markers of fat body cell clusters identified by FindAllMarkers.

Supplementary file 4: Top 20 most significantly expressed genes by cluster for ontology visualization.

Supplementary file 5: Proportion of cell cluster expressing selected immune and metabolic genes.

Supplementary file 6: Metabolites detected from midgut and fat body tissues at 2 and 7 dpbm.

